# Actin-inspired feedback couples speed and persistence in a Cellular Potts Model of cell migration

**DOI:** 10.1101/338459

**Authors:** Inge M. N. Wortel, Ioana Niculescu, P. Martijn Kolijn, Nir Gov, Rob J. de Boer, Johannes Textor

## Abstract

Cell migration is astoundingly diverse. Molecular signatures, cell-cell and cell-matrix interactions, and environmental structures each play their part in shaping cell motion, yielding numerous different cell morphologies and migration modes. Nevertheless, in recent years, a simple unifying law was found to describe cell migration across many different cell types and contexts: faster cells turn less frequently. Given this universal coupling between speed and persistence (UCSP), from a modelling perspective it is important to know whether computational models of cell migration capture this speed-persistence link. Here, we present an in-depth characterisation of an existing Cellular Potts Model (CPM). We first show that this model robustly reproduces the UCSP without having been designed for this task. Instead, we show that this fundamental law of migration emerges spontaneously through a crosstalk of intracellular mechanisms, cell shape, and environmental constraints, resembling the dynamic nature of cell migration *in vivo*. Our model also reveals how cell shape dynamics can further constrain cell motility by limiting both the speed and persistence a cell can reach, and how a rigid environment such as the skin can restrict cell motility even further. Our results further validate the CPM as a model of cell migration, and shed new light on the speed-persistence coupling that has emerged as a fundamental property of migrating cells.

**SIGNIFICANCE:** The universal coupling between speed and persistence (UCSP) is the first general quantitative law describing motility patterns across the versatile spectrum of migrating cells. Here, we show – for the first time – that this migration law emerges spontaneously in an existing, highly popular computational model of cell migration. Studying the UCSP in entirely different model frameworks, *not* explicitly built with this law in mind, can help uncover how intracellular dynamics, cell shape, and environment interact to produce the diverse motility patterns observed in migrating cells.

## INTRODUCTION

Imagine a T cell moving in the outer layer of the skin. Tasked with patrolling the epidermis, it scans for early signs of re-invasion by pathogens known from earlier attacks. Its movement is rapid yet undirected: with narrow protrusions almost resembling dendrites, it probes its surroundings before choosing where to go next. Its decision made, it squeezes its way through the tight junctions between the skin’s keratinocytes – moving its attention to unexplored areas (1). Suddenly, the scene changes as a cut disrupts the tissue layer. The released damage signals attract neutrophils, which rapidly crawl towards the wound with a motion far more directed than that of the T cell patrolling this site earlier. Upon arrival, they stimulate the movement of yet another cell population: the epithelial sheet adopts a directed, slow-but-steady collective motion that (combined with proliferation) eventually closes the wound. Homeostasis is restored (2, 3).

The above scenario illustrates just a few of the many movement patterns and phenotypes found among migrating cells. While all mammalian cells share the same basic mechanism of actomyosin-driven cell motion, differences in their molecular signatures – as well as in the structure of the environment they move in – nevertheless produce a rich spectrum of different migration modes (4). Movement can be fast or slow, in a persistent or frequently changing direction, across a 2D surface or inside a 3D matrix. Cells can be round or elongated, forming narrow or broad protrusions that do or do not rely on focal adhesions. Cells can move as isolated individuals, or collectively as a cohesive sheet or stream.

Interestingly – despite these highly diverse migratory behaviors observed in different cell types and contexts – one universal law seems to describe motion patterns across many migrating cells in various controlled experimental settings: faster cells move more persistently (5–7). Maiuri et al proposed that this Universal Coupling between Speed and Persistence (UCSP) arises from a positive feedback on cell polarity mediated by the actin cytoskeleton. Because specialized clutch molecules provide friction between the actin filaments and the cell’s surroundings, these actin filaments move backwards in the reference frame of the moving cell. This “actin retrograde flow” depends linearly on cell speed. A theoretical model revealed how actin retrograde flow can also stabilize cell polarity (and thus persistence) if it transports polarity cues towards the cell’s rear end. The resulting polarity cue gradient in turn stabilizes actin retrograde flow in a positive feedback loop. Thus, higher speeds are linked to higher persistence because they stabilize cell polarity via the actin retrograde flow (7). As actin retrograde flow is a highly conserved feature of cell migration, the UCSP holds for cell types with very different migration modes.

Maiuri et al proposed this theoretical model to explain how the UCSP arises from the actin-based advection mechanism described above (7). However, the apparent universality of this migration law raises the question to what extent other, *existing* models of cell migration might capture this phenomenon. If the UCSP is truly so universal, it might be important to include it in other models used to study cell migration patterns. And vice versa, studying the UCSP in different models may shed new light on this fundamental law of cell migration.

Many mathematical and computational models of cell migration already exist. The complex interplay between cell-intrinsic and environmental factors has made cell migration a popular topic of study among computational biologists, raising questions such as: How do single cells coordinate their motion to start moving collectively, as a single whole? How does each cell’s molecular machinery interact with its shape during motion? How do constraints on cell shape posed by a crowded tissue environment alter the immune cell migration patterns we see? And does it matter? What migratory pattern *should* immune cells adopt to find their targets most efficiently? Over the years, many studies have shed light on these questions by studying the movement of both single cells and cell collectives with a combination of experimental and modelling approaches (8–12).

The models used are diverse, ranging from relatively simple particle models to detailed physical models linking intracellular signalling and cell shape (12, 13). These models differ in how they describe motion: while motion characteristics such as speed and persistence are *input* parameters in the simpler particle models, in the more complex mechanistic models, motion emerges as *output* of the underlying dynamics (which can involve a combination of molecular signalling, cell shape, and environmental factors, depending on the models used). While a correlation between speed and persistence can simply be imposed by the user in the former case (14), this is impossible in the latter case, where speed and persistence are emergent outcomes outside of the user’s control. A more empirical approach is therefore needed to examine whether such models can also capture a phenomenon like the UCSP.

Here, we therefore examine such a model in which speed and persistence emerge from a migration machinery that interacts with the cell’s shape and the environment. We focus on the Cellular Potts Model (CPM), a popular framework for modelling cell migration because it naturally captures complex cell shapes and cell-environment interactions (1, 10, 15). Rather than developing a new model, we explore an existing model known to reproduce cell migration with realistic cell shapes: the “Act-CPM” (15). Importantly, its spatial nature also allowed us to investigate how this cell-intrinsic migration machinery interacts with more complex tissue environments. While this particular model was not built with the UCSP in mind, we ask whether it nevertheless captures this seemingly fundamental law – and how this is influenced by the environment. The original experimental data mostly established the UCSP in simple, obstacle-free geometries (straight 1D adhesive lines or a free 2D surface) (5–7). This raises the question whether the UCSP remains a universal law across the more diverse environments cells face *in vivo*.

Interestingly, we find that in our existing CPM model, the UCSP emerges spontaneously in simulations of cell migration in various environments. We also show that studying the UCSP in this model yields novel insights: the spatial nature of the Act-CPM allowed us to examine how cell shape dynamics interact with the UCSP to determine the migratory patterns a cell can adopt. Finally, we also show that the UCSP may not be universal throughout *all* environments, because a strongly restrictive environment may overrule speed-persistence coupling in T cells patrolling the epidermis.

## METHODS

### Model

Our model is an extension of the Cellular Potts Model (CPM), which represents cells as collections of pixels on a 2D or 3D grid. Essentially, these cells “move” by randomly trying to add or remove pixels at their borders in “copy attempts” (Figure 1). While doing so, cells try to minimize the energetic cost Δ*H* associated with maintaining their shape and contacts with neighbor cells (Figure 1A). Thus, this model is spatial since each pixel corresponds to a region in space. We get temporal dynamics by performing several copy attempts in a row; time is then measured in Monte Carlo Steps (MCS), where every MCS, we perform a number of copy attempts equal to the number of pixels on the grid.

**Figure 1:**
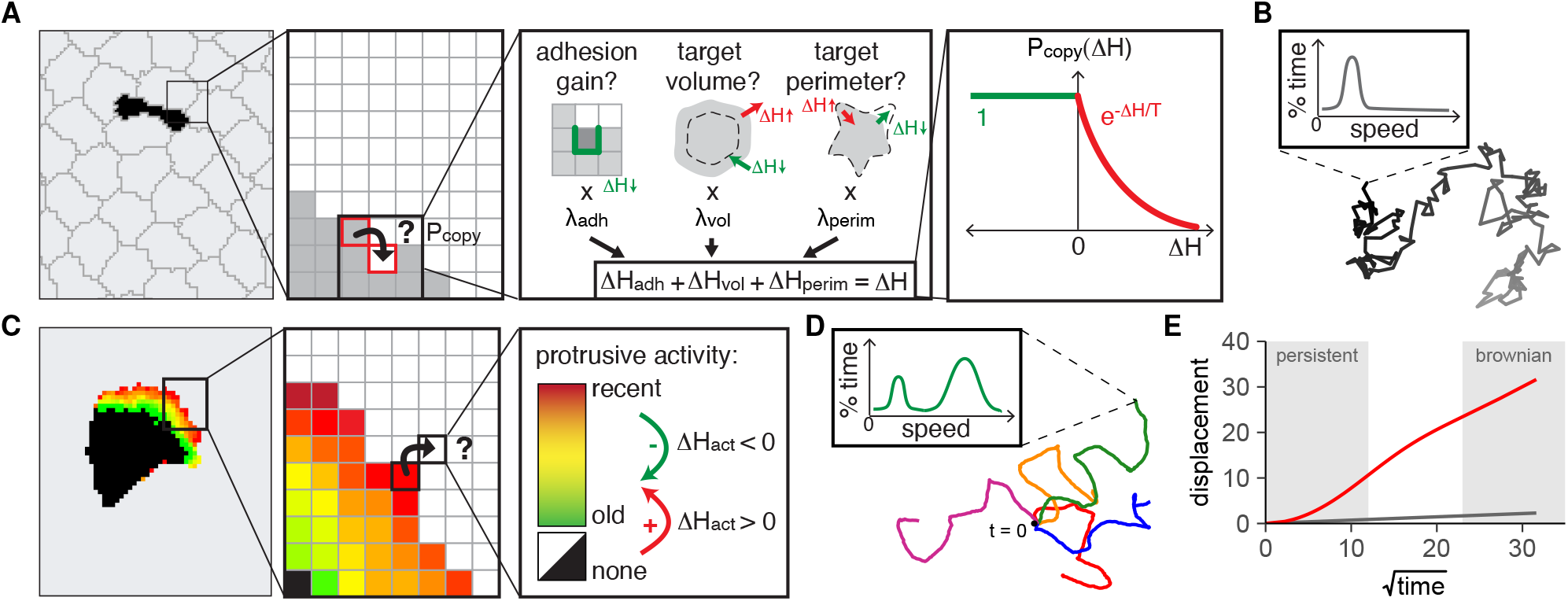
*In silico* simulation of shape-driven cell migration within complex environments. **(A)** A CPM represents a tissue as a collection of pixels on a grid, each belonging to a specific cell or the extracellular space. Pixels randomly try to copy their cell identity into pixels of neighbor cells, with a success probability P_copy_ depending on how that change would affect the “physical” properties of the involved cells (cell-cell adhesion, and deviation from target volume and/or perimeter, dashed lines). The weighted sum of these energetic effects (ΔH) is negative for energetically favourable copy attempts. **(B)** In a CPM with only adhesion, volume, and perimeter constraints, cells only exhibit Brownian, diffusion-like motion. Plot shows an example cell track. The inset illustrates that the distribution of instantaneous speeds remains small throughout the track. **(C)** In the Act-CPM (15), each pixel’s “activity” represents the time elapsed since its most recent successful protrusion. Copy attempts from active to less active pixels are stimulated (negative ΔH_act_), whereas copy attempts from inactive to more active pixels are punished (positive ΔH_act_). **(D)** Act cells can alternate between persistent motion and “stops” in which they change direction (intermittent random walk, “I-RW”). Plot shows example tracks of 5 Act cells with overlaying starting point (black dot, t = 0). The inset illustrates the distribution of instantaneous speeds for this I-RW, “stop-and-go” motion, which has peaks at zero (representing the “stops” in the track) and at high speeds (representing the “go” intervals). **(E)** Displacement plot of CPM cells. Brownian motion (without the Act extension, gray line) results in a linear curve. Act cell movement appears as Brownian motion on large time scales (linear part of red line), but is persistent on smaller time scales (non-linear start of red line).

The above-mentioned tendency to minimize the global energy arises because we do not let all copy attempts succeed; rather, the success rate of each attempted copy depends on its energetic effect as follows:

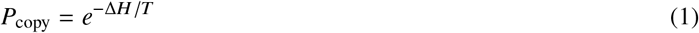

where the temperature *T* controls the amount of randomness in the simulation. Here, a copy attempt’s success chance depends on its “energetic cost” Δ*H*, which in turn depends on how that change would affect the global energy or *Hamiltonian H* of the system. We define this energy *H* in terms of energetic “constraints”, which may vary between different CPM models but often include cell-cell adhesion and constraints on cell volume and perimeter (as we will do here). Since all cells contribute to the total energy *H*, *P*_copy_ depends not only on the shape of the cell trying to move, but also on that of the cell it tries to displace.

This general framework allows CPMs to reproduce realistic, dynamic cell shapes and -behavior using only a few simple rules and parameters. Their spatially explicit yet simple nature makes CPMs powerful tools for modelling the interactions of individual cells with complex, multicellular environments in a controlled setting. However, the energy that basic CPM cells try to minimize is based solely on adhesion and cell shape. As there is no energetic benefit for consistent movement in any given direction, cells undergo a Brownian, diffusion-like motion rather than actual migration (Figure 1B).

We therefore use an extension of the CPM that does allow for active migration (15). In this “Act-CPM”, pixels newly added to a cell temporarily remember their recent protrusive activity for a time of max_act_ MCS (where max_act_ is a model parameter). Copy attempts from active into less active pixels are rewarded via a negative energy contribution Δ*H*_act_ to the overall energy cost Δ*H*, stimulating copy attempts from a source pixel *s* in an “active” region into a less active target pixel *t*:

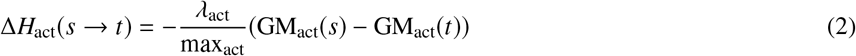

where GM_act_(*p*) of pixel *p* is the geometric mean of the activity values of all pixels in the (Moore) neighborhood of *p*. The division by max_act_ ensures a normalization; what matters is the *relative* activity, not its absolute value. Thus, max_act_ controls the steepness and temporal stability of the activity gradient, whereas the Lagrange multiplier *λ*_act_ determines its contribution to the total energy Δ*H* associated with the copy attempt. Δ*H*_act_ is negative when GM_act_ (*s*) > GM_act_(*t*).

By adding this term to the CPM, we obtain a positive feedback loop wherein recently added pixels become more likely to protrude again (Figure 1C). Consequently, local groups of active pixels form stable protrusions that drag the cell forward in persistent motion before disappearing again, at which point the cell stops until a new protrusion forms. Thus, cells alternate between intervals of persistent movement and “stops” where they can switch direction (Figure 1D), a pattern also called an “intermittent random walk” (I-RW) (7). On larger time scales, movement still resembles Brownian motion (new protrusions form in random directions). Persistence is only evident on the smaller time scales during which the cell has a stable protrusion and maintains its direction of movement (Figure 1E).

This I-RW behavior qualitatively resembles the characteristic “stop-and-go” motility of T cells searching for antigen in the lymph node (16, 17), as well as the motility of other cell types (7). Because this model can also simulate cell interactions and migration within tissues (15), this makes it a suitable tool for examining the effect of tissue context on cell migration *in silico*.

All simulations were built using Artistoo (18). For details on parametrisation and exact setup, see the Supplementary Methods. All simulation and analysis code will be made available at http://github.com/ingewortel/2020-ucsp upon publication of this manuscript. For a complete description of the CPM, see (15).

### Analysis

During each simulation (every 5 MCS), we recorded both the position of the cell’s centroid (to compute speed and persistence time), and several other cell properties (to keep track of the cell’s shape and degree of polarization).

#### Quality control: shape and polarization

In the CPM, the pixels belonging to a single cell are held together mostly via the adhesion term in the Hamiltonian, which favours cell shapes where pixels belonging to the same cell adhere to each other. However, this adhesive force can become negligible relative to the other Δ*H* terms – for example when Δ*H*_act_ is large due to a high *λ*_act_. Thus – especially in 3D – cells may break apart at high values of *λ*_act_, despite the unfavourable changes in adhesion energy associated with this break.

As frequent cell breaking causes artefacts in the tracking data that may bias the measurement of speed and persistence, it is important to use parameter ranges that prevent such an unbalanced contribution of the different Δ*H* terms. To estimate the frequency of cell breaking, we therefore recorded the *connectedness* (*C*) of the cell every 5 MCS of each simulation. This number is 1 for an intact cell, and approaches 0 for a collection of unconnected pixels (see Supplementary Methods for details). For all simulations reported in this manuscript, we checked that a connectedness below 0.95 did not occur in more than 5% of the measured values – ensuring that cells were intact for the majority of the simulation.

Additionally, as a measure of the total size of the active protrusion(s) of a cell, we counted the percentage of pixels of that cell with an Act-CPM activity > 0. Ranges of max_act_ and λact were chosen such that protrusions made up > 0 but < 100% of the cell’s total volume.

#### Track analysis: speed and persistence time

Cell centroids were recorded at regular time intervals (5 MCS) to reconstruct cell trajectories or “tracks”. All simulated cell tracks were analyzed in R (version 3.6.1) using the celltrackR package (version 0.3.1) (19) to compute speed and persistence time. Speeds were computed from instantaneous “step” speeds along the track, and persistence time was defined as the half life of the autocovariance curve. Analyses were performed in a step-based manner (combining steps from independent tracks for robustness), using separate groups of 5 tracks each to estimate variation. See Supplementary Methods for details.

## RESULTS

### The Act-CPM reproduces the UCSP observed in migrating cells

Given the apparent universality of the UCSP, we first tested whether our CPM could reproduce the phenomenon as observed in experimental data. Among cell tracks recorded under standardized conditions, Maiuri et al. (5, 7) saw that all cell types analyzed followed one general rule: faster cells turned less frequently. Most of these experiments involved the migration of cells moving along adhesive tracks or within microchannels (5, 7). To mimic this “one-dimensional” experimental set-up *in silico*, we constrained Act cells between two parallel walls, leaving a space of 10 pixels within the channel (Figure 2A). This set-up resulted in cell elongation comparable to that observed for cells moving on 1D adhesive tracks (compare Figure 2A to Figure 1B in (7)). Act cells moving in these microchannels reproduce I-RW behavior, migrating persistently in one direction until they lose their active protrusion – at which point they wait for a new protrusion to form and can stochastically switch direction (Figure S1A and Movie S1, Interactive Simulation S1 in Supplementary Materials).

**Figure 2:**
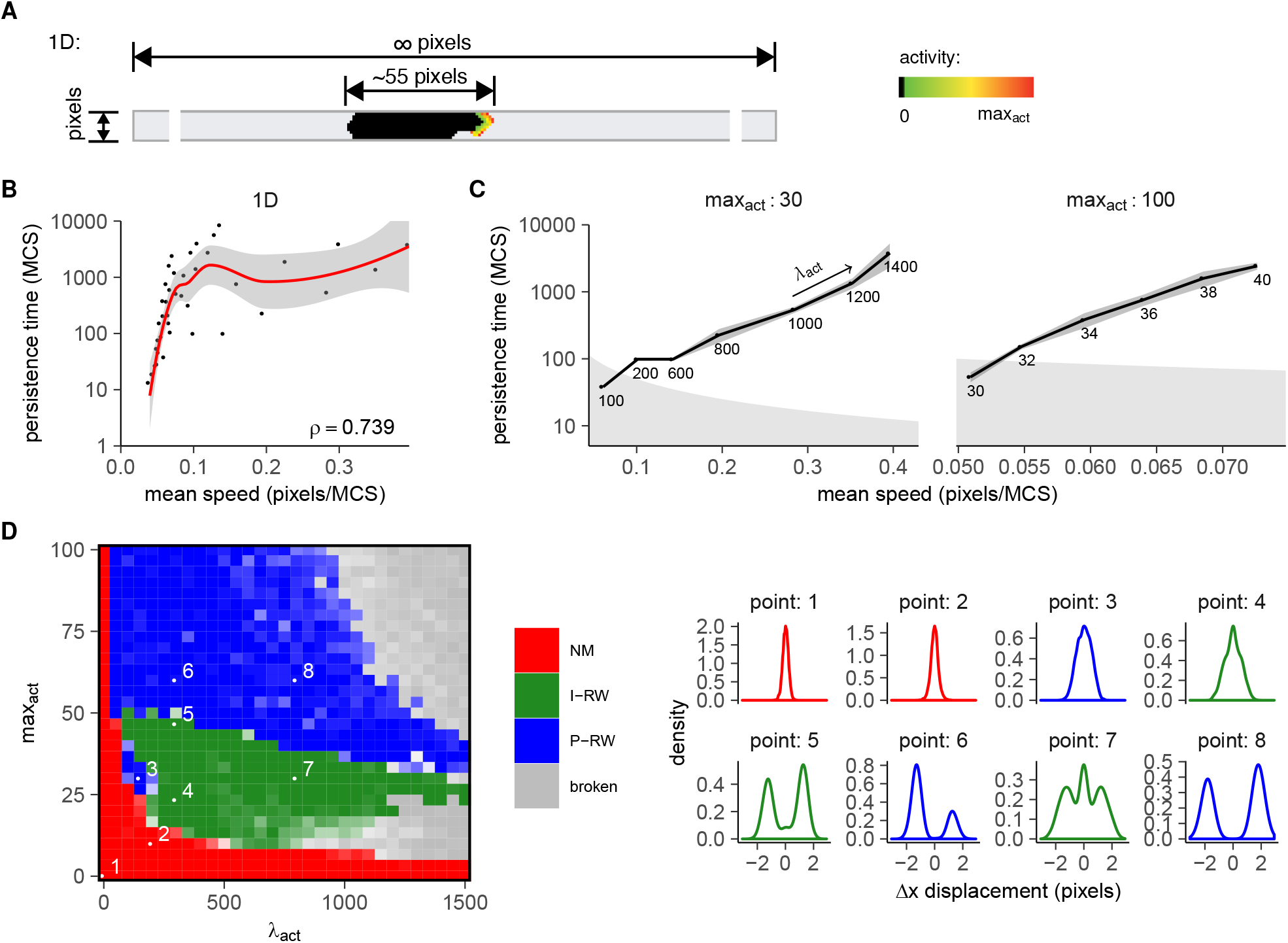
The Act-CPM reproduces the UCSP observed in experimental data. **(A)** Set-up and example Act cells for simulations of “1D” migration in microchannels. Color gradients indicate the active protrusion. Simulated microchannels consist of two parallel walls, leaving 10 pixels in between. **(B)** An “exponential” speed-persistence coupling arises in the Act-CPM (i.e., there is a range in which speed is proportional to the logarithm of persistence time; ρ = spearman correlation coefficient). Both max_act_ and *λ*_act_ were varied; see Supplementary Methods for a list of parameters used and (C) for the relationship at fixed max_act_. **(C)** Speed-persistence coupling is stronger for cells with the same max_act_. Plot shows mean ± SD of persistence time plotted against speed, for two values of max_act_; numbers in the plot indicate the corresponding value of *λ*_act_. Shaded gray areas in the background indicate regions where the persistence time is lower than the time it takes for the cell to move 10% of its length. **(D)** Phase diagram of migration modes in microchannels for different max_act_ and *λ*_act_ (left), as based on the displacement distributions (right). Cells were classified as non-migratory (NM) if they hardly moved (displacement distribution with a single peak centered at 0). Cells were classified as P-RW if the displacement distribution had two clear peaks (for motion to the left and right, respectively) and as I-RW if it had three peaks (with an extra peak at zero for the “stops”); see Supplementary Methods for details. This classification yielded fairly consistent “phases” in the parameter space, although it was harder to distinguish peaks for cells that were barely moving (e.g. point 3). Some *λ*_act_ and max_act_ combinations in the CPM are not viable because the protrusion tear the cell apart; these were classified as “broken”. In the diagram, colors represent migration mode, whereas the intensity of the color represents agreement of the classification between different independent estimates (see Supplementary Methods).

We then used this set-up to simulate cells with different migratory behavior. Two parameters control migration in the Act-CPM. The first, *λ*_act_, tunes the contribution of the positive feedback Δ*H*_act_ relative to the other energetic constraints (adhesion, volume, perimeter), and can be interpreted as the force exerted on the cell membrane by actin polymerization at the cell’s leading edge. When *λ*_act_ is large, this force can easily push the membrane forward to form a stable protrusion, but when it is small, the actin cytoskeleton has a hard time overcoming other, opposing forces (such as membrane tension). Higher *λ*_act_ values therefore yield larger protrusions (Figure S1B in Supplementary Materials). The second parameter, max_act_, determines how long pixels remember their activity (measured in MCS, the time unit of the CPM). It thereby limits the protrusion width (i.e., how much it extends into the cell interior), and can be interpreted as the lifetime of polymerized actin. Higher max_act_ values therefore have a stabilizing effect on the protrusions – yielding larger protrusions even at small forces *λ*_act_ (Figure S1B in Supplementary Materials).

To examine the UCSP across cells with different migratory behaviors, we generated cell tracks for Act cells with various *λ*_act_ and max_act_ values, and subsequently extracted the speed and persistence time from those tracks. (Here, the “persistence time” is the average time over which the direction of motion changes, computed from the speed autocorrelation function as described in Supplementary Methods). This analysis revealed a weak exponential coupling between speed and persistence (Figure 2B). Although the correlation was weak in this heterogeneous dataset of Act cells with highly different *λ*_act_ and max_act_ parameters, it became much stronger when we stratified cells by max_act_ value (Figure 2C). Analysis of speed and persistence in these tracks yielded the same exponential correlation between speed and persistence as was observed in experimental data (7). This finding was independent of the choice of max_act_, as we found similar curves for both values of max_act_ (Figure 2C and see below).

Although changes in max_act_ did not affect speed-persistence coupling, they were associated with a change in migration mode. Whereas cells with lower max_act_ values switched from a non-migratory (NM) phenotype to I-RW “stop-and-go” motion as *λ*_act_ increased, cells with higher max_act_ values instead went from non-migratory to a “persistent random walk” (P-RW) mode with hardly any stops. To illustrate this difference, we computed a “phase diagram” of migration behavior with varying max_act_ and *λ*_act_, using the distribution of displacements to determine migration modes (Figure 2D). This phase diagram strongly resembled that obtained by Maiuri et al using the original UCSP model (7) (See Appendix A in Supplementary Materials for a more extensive comparison between the two models).

The exponential speed-persistence relationship is mediated by the *λ*_act_ parameter: whereas speed increased linearly with *λ*_act_ (Figure S2A), persistence time increased exponentially at higher *λ*_act_ values (Figure S2B). Indeed, the linear relationship between the Langrange multiplier λ and cell speed has previously been explained for a similar CPM migration model based on a chemotactic, rather than cell-intrinsic, force (20). In addition, we find that the exponential relationship between *λ*_act_ and persistence time follows directly from the kinetics of the CPM and from the size of the “energy barrier” cells need to cross to lose an active protrusion (Appendix B in Supplementary Materials). Thus, *λ*_act_ exponentially couples speed and persistence in this CPM model of cell migration. These results demonstrate that the Act-CPM reproduces the UCSP observed in migrating cells, as well as the underlying different migration modes.

### Speed-persistence coupling in the Act-CPM spans a range of migration modes

We then exploited the spatial nature of the Act-CPM to examine the UCSP in environments less artificial than a microchannel, allowing cells to take on their preferred shapes (Figure 3).

**Figure 3:**
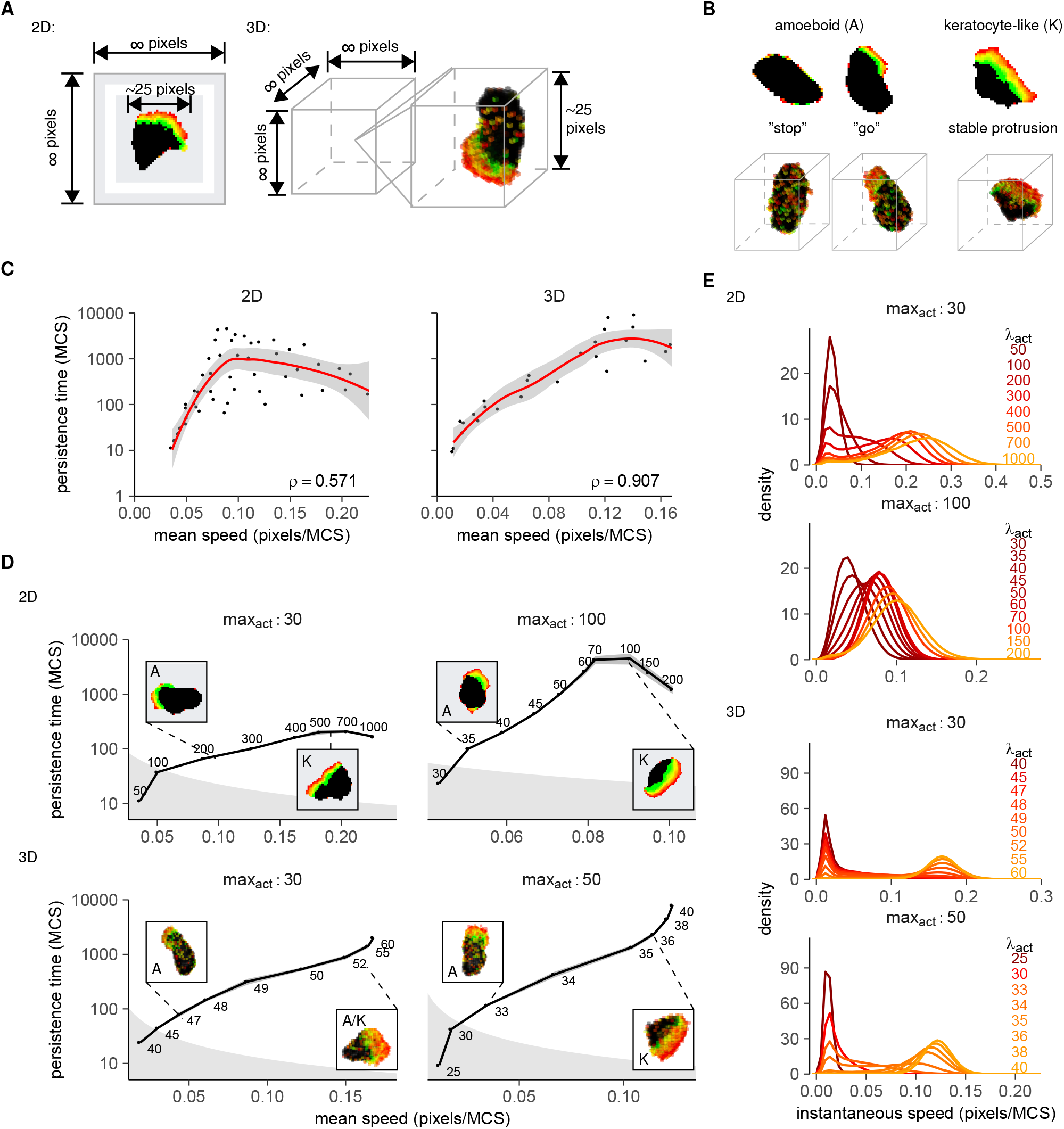
Speed-persistence coupling spans a range of migration modes. **(A)** 2D & 3D simulations were performed in an empty grid. **(B)** Migration modes in the Act-CPM; see also (15). Amoeboid cells form small, narrow protrusions that decay quickly and produce stop-and-go motion. Keratocyte-like cells have broader, more stable protrusions. **(C)** Exponential speed-persistence coupling in 2D and 3D for different (*λ*_act_, max_act_). ρ = spearman correlation coefficient). See Supplementary Methods for a list of parameters used. **(D)** Speed-persistence coupling in 2D and 3D becomes stronger for cells with the same max_act_ and spans a transition from amoeboid to keratocyte-like motion. Plots show mean±SD persistence time plotted against speed. Insets show representative cell shapes; regions are shaded gray where the persistence time is below the time needed to move 0.1 cell length. See also Figure S3A. **(E)** Instantaneous speed distributions of 2D and 3D Act cells. Cells transit from not moving (single peak at speed ~0 pixels/MCS), via “stop-and-go” motility (bimodal distributions), to near-continuous movement (single peak at high speed). See also Figure S3B.

In addition to discovering the UCSP in cells navigating “1D” adhesive tracks, Maiuri et al. (7) also confirmed this coupling in cells migrating on surfaces (“2D”) and within 3D environments. We mimicked these experiments by simulating Act cell migration in large, unconfined 2D and 3D spaces (Figure 3A). Like in the microchannel data (Figure 2B,C), we again found a weak exponential correlation between speed and persistence (Figure 3C) that became stronger when we stratified cells by max_act_ value (Figure 3D, Figure S3A in Supplementary Materials). In fact, the exponential increase in persistence was now accompanied by a transition in cell shapes (insets in Figure 3D).

An important feature of the Act-CPM is that it reproduces different cell shapes and migration modes (15). In contrast to the uniform, elongated shape observed in channels, Act cells moving in 2D and 3D can form different types of protrusions (15) (Figure 3A,B,D and Movie S2, Interactive Simulation S2, S3 in Supplementary Materials). Low values of *λ*_act_ and max_act_ promote the formation of small and narrow protrusions that form and decay dynamically, giving rise to an amoeboid (“stop-and-go”, I-RW) migration mode (Figure 3B, left, Figure 3D, and Movie S2 in Supplementary Materials). By contrast, large values of *λ*_act_ and/or max_act_ favor the formation of broad, stable protrusions, yielding a more persistent “keratocyte-like” (P-RW) migration mode (Figure 3B, right, Figure 3D, and Movie S2 in Supplementary Materials). This transition occurred in both the 2D and the 3D model, although we note that the “amoeboid” behavior in 3D was slightly different in 3D than in 2D. In 3D, the “stops” tended to be longer, and many protrusions were too unstable to make the cell move far from its place. The “go” intervals were less frequent and required the cell to take on a somewhat broadened shape, although not as broad as in the 3D “keratocyte-like” motion (Movie S2, Interactive Simulations S2 and S3 in Supplementary Materials).

This transition between different migration modes is illustrated by the distributions of the instantaneous speeds from our simulated cell tracks (Figure 3E, Figure S3B in Supplementary Materials). The bimodal shape of this distribution – especially evident at low values of max_act_ – reflects the “stop-and-go” I-RW behavior of migrating Act cells: when the cell is in a “stop”, it has a very low instantaneous speed of almost 0 pixels/MCS, whereas the “go” intervals of movement are responsible for the peaks at a higher speed (This is similar to the displacement distributions used for the phase diagram in Figure 2D, except now we have more than one dimension and and infinite number of directions rather than just two. We therefore look only at the *magnitude* of the displacement/velocity vector – so the separate peaks for “left” and “right” motion become one.). Thus, at very low *λ*_act_, the cell barely moves at all – as indicated by a single peak at instantaneous speeds of almost zero (Figure 3E). This corresponds to a non-migratory (NM) cell without protrusions, spending most of its time in “stops”. As *λ*_act_ increases, the cell enters the “stop-and-go” (I-RW), amoeboid migration regime (bimodal distributions) Higher *λ*_act_ values not only increase migration speed (reflected by an upward shift of this second peak), but also reduce the amount of time a cell spends in “stops” (reflected by a decrease in the size of the first peak). As stops provide an opportunity for the cell to change its direction (Figure 3B), the reduced “stopping time” at high *λ*_act_ values explains why Act cells with high *λ*_act_ values migrate not only faster, but also more persistently. Finally, at the highest max_act_ and *λ*_act_ values, the cell takes on a keratocyte-like shape and almost never stops moving (P-RW).

Together, these results demonstrate that the exponential speed-persistence coupling holds in 2D and 3D and spans different “regimes” of migration.

### Both Act cell speed and persistence saturate in a cell shape-dependent manner

Interestingly, our data also show the saturation of the persistence at higher cell speeds that was reported in the experimental data (compare the 2D figures in Figure 3D to the data in (7)). In fact, this saturation was not limited to persistence. Whereas speed initially increased linearly with *λ*_act_, it plateaued at higher *λ*_act_ values (Figure 4A,B, Figure S3C in Supplementary Materials). The maximum speed reached depended on the protrusion shape-parameter max_act_ (Figure S3D,E in Supplementary Materials), and in all cases, the initial linear part of the graph spanned the entire transition from amoeboid to a keratocyte-like shape. This finding suggests that having to maintain a broad protrusion limits the speed a cell can reach. In line with this idea, we did not observe this saturation in microchannels, which prevent the cell from acquiring the broad protrusions observed in 2D and 3D (Figure S2A in Supplementary Materials).

**Figure 4:**
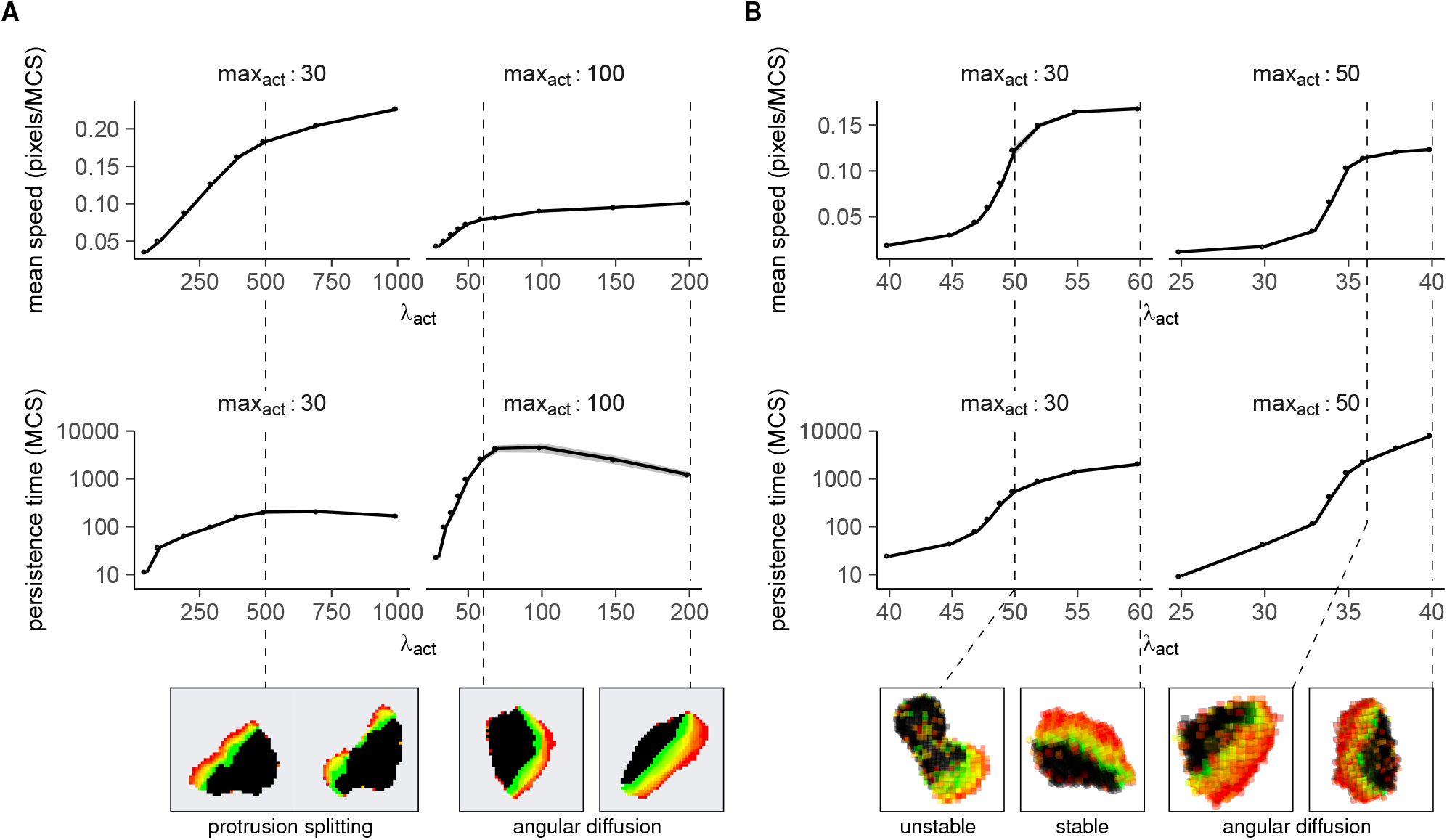
Cell shape dynamics limit both the speed and persistence of migrating cells. Mean ± SD of speed and persistence time of **(A)** 2D and **(B)** 3D Act cells, plotted against *λ*_act_ for different values of max_act_. Open circles indicate points where the persistence time is lower than the time it takes the cell to move 10% of its length (corresponding to the points in the gray background in Figure 3B). Insets show cell shapes at the indicated parameter values.

Similarly, the cell shape changes observed in 2D and 3D seem to put an upper bound on persistence that disappears when the cell is constrained by a microchannel (Figure 4A,B, Figure S3C-E, S2B in Supplementary Materials). The initial exponential increase in persistence again spanned the entire transition from amoeboid to keratocyte protrusion shapes, before eventually saturating at a max_act_-dependent value. Again, this phenomenon appears to be linked to protrusion shapes. Whereas cells with low max_act_ do tend to form keratocyte-like protrusions at high *λ*_act_ values, these protrusions do not extend far into the cell and are prone to splitting – forcing the cell to turn towards one of the protrusion halves (Figure 4A and Movie S3 in Supplementary Materials). Although higher max_act_ values allow for larger persistence times by letting broad protrusions extend farther into the cell and preventing them from splitting (Figure 4A,B), persistence still saturates eventually due to slight, stochastic turning of the stable protrusion around the cell perimeter (“angular diffusion”, Figure 4A,B and Movie S3 in Supplementary Materials) (7).

By showing how the shape of migrating cells puts a natural upper bound on both the speed and the persistence a cell can reach, these results explain the saturation of persistence observed by Maiuri et al (7). However, there was a striking effect of dimensionality on this process: although we observed shape-driven saturation in both 2D and 3D, the shape of the speed-persistence curve was different for 2D and 3D simulations (Figure 3B, 4). In both settings, speed and persistence saturated at high *λ*_act_ after an initial increase (which was linear for speed and exponential for persistence). Yet, whereas persistence saturated before speed in 2D (Figure 4A), 3D Act cells showed a much stronger saturation of speed that preceded the saturation of persistence (Figure 4B). Thus, when both speed and persistence have a natural upper bound, the dominant saturation effect can be context-dependent – altering the shape of the speed-persistence curve.

### Environmental constraints break the UCSP for T cell migration in the epidermis

So far, our models mimicked the environments in which the UCSP was initially discovered, where cells can migrate rather easily. But many cell types also need to move in crowded or stiff environments that strongly constrain cell shape. To investigate how such constraints would impact speed-persistence coupling, we modelled T cell migration in the epidermal layer of the skin. As one of the key entry points through which pathogens can enter the body, healthy skin contains substantial numbers of T cells (21). T cells attracted to the epidermis during an infection can remain there for a long time: even a year after the resolution of an infection, specific T cells still persist in the same region of the epidermis (22–25). Whereas subtle chemotaxis guides T cells towards infected cells during the effector phase (26), these remaining T cells actively patrol the epidermis without such chemotactic guidance (1) – migrating in patterns shaped by a combination of cell-intrinsic factors and environmental constraints. Importantly, even though the tight contacts between keratinocytes make the epidermis one of the most rigid environments T cells encounter *in vivo*, T cells in the epidermis are nevertheless highly motile (1).

We therefore focused on this extreme example to examine how environmental structure can affect the UCSP. To this end, we simulated T cell migration in the skin as reported previously (15), placing an Act cell in a grid covered completely with keratinocytes (Figure 5A). In this setting, Act T cells move by squeezing in between the keratinocytes (Movie S4) – but because of the opposing forces from the surrounding keratinocytes, cells now required higher *λ*_act_ forces to counter this resistance and start moving (Figure S4 in Supplementary Materials). At sufficiently high *λ*_act_ values, they once again showed the characteristic “stop-and-go” motility before eventually switching to near-constant motion with hardly any stops (Figure 5B, Figure S4, Movie S4, and Interactive Simulation S4 in Supplementary Materials).

**Figure 5:**
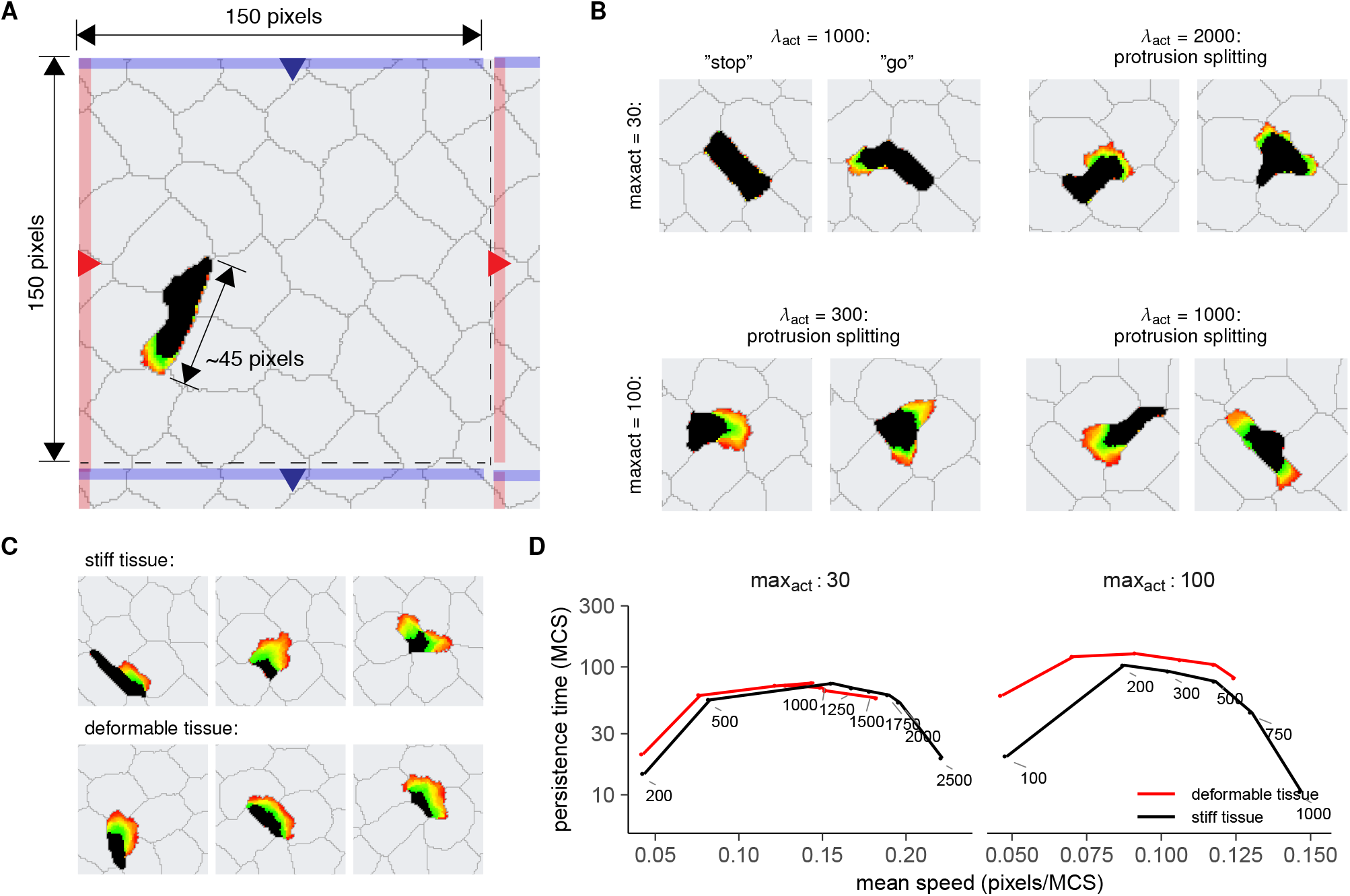
Environmental constraints limit T cell persistence in a model of the epidermis. **(A)** An Act T cell (black) moving in between keratinocytes (gray) in the epidermis. Simulations were performed in a 150 × 150 pixel grid with linked borders (for example, a cell moving off the grid towards the red region on the right re-enters the grid at the equivalent red region on the left). **(B)** Shapes of Act T cells constrained between keratinocytes. At lower *λ*_act_/max_act_ values, T cells show typical amoeboid “stop-and-go” behavior. At higher *λ*_act_ and/or max_act_ values, cells do not obtain a broad, keratocyte-like shape like they normally would (Figure 3), but stay elongated due to environmental constraints. At junctions between keratinocytes, however, protrusions tend to split. **(C)** Whereas formation of broad protrusions is mostly prevented in “stiff” skin tissue, Act cells in a more deformable tissue can form broad protrusions by pushing apart surrounding cells. **(D)** Mean persistence time plotted against speed for different combinations of *λ*_act_ and max_act_, tissues with different stiffness. Shaded gray background indicates regions where the persistence time is lower than the time it takes for a cell to move 10% of its length.

Unlike Act cells in an unconstrained environment, these Act T cells could not fully switch from amoeboid to keratocyte-like cell shapes as *λ*_act_ and/or max_act_ increased (Movie S4). Even though cells at high *λ*_act_/max_act_ became somewhat broader, they still mostly maintained their amoeboid shape, probing their surroundings with narrow protrusions and migrating in the direction of their longest axis. However, when these cells approached a “T-junction”, they sometimes formed a broad protrusion in the space between the keratinocytes that eventually split up into two separate protrusions going in opposite directions (Figure 5B). This protrusion splitting caused the cell to slow down until one of the two active regions gained the upper hand (Movie S4).

In this set-up, increases in *λ*_act_ were once again associated with a higher speed that gradually saturated at high *λ*_act_ values (Figure S5A in Supplementary Materials), but persistence times now saturated much earlier, reaching a plateau at ~90 MCS for max_act_ = 30 and 140 MCS for max_act_ = 100 (Figure S5B in Supplementary Materials). With cell speeds around ~0.12 and ~0.07 pixels/MCS, respectively, this corresponds to persistent movement over distances in the range of ~10-12 pixels – just under the distance the T cell can travel before arriving at another junction (Figure 5A, Figure S5C in Supplementary Materials).

Thus, the structure of the environment appears to be a limiting factor for T cell persistence in this scenario. Indeed, when we placed cells in a grid covered with more deformable cells, cells with a high max_act_ of 100 could once again form their preferred broad protrusions and move in straighter lines by pushing the surrounding cells apart (Movie S5, Figure 5C). This in turn resulted in a higher persistence (Figure S5B) and a slighly increased speed (Figure S5A). By contrast, cells with a low max_act_ of 30 – which cannot stably form broad protrusions even when unconstrained by tissue (Figure 4A) – had similar speed and persistence in the limit of high *λ*_act_ values, regardless of tissue stiffness (Figure S5A,B). These results demonstrate how interactions between the environment and cell shape determine how strongly the tissue affects motility patterns.

The observed rapid saturation of persistence eclipsed the UCSP for T cells migrating in the skin, removing the speed-persistence correlation (Figure 5D). This result was independent of the rigidity of the surrounding keratinocytes: although a reduction in tissue stiffness slightly increased the maximum persistence time for cells with high max_act_ (Figure S5B in Supplementary Materials), this did not rescue speed-persistence coupling (Figure 5D). Thus, although the UCSP appears to be valid for all migrating cells, cell-intrinsic speed-persistence coupling may be obscured when environmental factors place additional, more stringent constraints on persistence.

## DISCUSSION

The UCSP is a simple yet highly general quantitative law of cell motion, which holds across a broad spectrum of migrating cells. Given the incredible diversity of the mechanisms driving cell migration, it is remarkable that such a general law exists at all. Nevertheless, after the UCSP’s initial discovery (5), Wu et al. (6) later also found a robust speed-persistence coupling in an independent study. Maiuri et al. (7) explained it by showing that actin retrograde flow can mechanistically couple cell polarity to migration speed, and Yolland et al. (27) further strengthened this explanation by demonstrating that the actin flow field controls stable cell directionality. Here, we confirm this seemingly fundamental law of cell migration in a completely different but popular modelling framework (the CPM), and show how cell shape dynamics and environmental constraints alter the shape of the speed-persistence correlation.

In the Act-CPM, migration arises through a combination of local activation (the Δ*H*_act_ force allowing protrusion) and global inhibition (the “membrane tension” that makes the cell retract its rear after the protruding front has stretched its perimeter beyond the target value). We find that this simple polarity mechanism not only suffices to reproduce the speed-persistence coupling observed in migrating cells (7), but also introduces an interaction between the UCSP and cell shape dynamics: the increase in speed and persistence is accompanied by a transition between different cell shapes and migration modes. Indeed, the combined local activation of protrusions and global inhibition via membrane tension dynamically link cell shape to cell motion in the CPM. Similar to shape-motility interactions observed in real cells (13), Act cell shapes are closely linked to both motility characteristics underlying the UCSP: speed and turning behavior.

The observed link between cell shape and speed is consistent with several other studies (9, 28–31). In T cells, this coupling arose from the same actin retrograde flow that also underlies the UCSP (7, 32). Lavi et al. (31) recently extended the original UCSP model to a 2D free boundary model with dynamic cell shapes, and again found that increased cell speeds correlated with broader cell shapes – similar to the shape changes observed in the Act-CPM for increasing *λ*_act_.

Likewise, cell shape is intricately linked to turning behavior in fish keratocytes. The broad protrusion (“lamellipodium”) of these cells normally allows them to migrate persistently. Yet, the stability of the lamellipodium partly depends on its shape, and shape changes of the cell may destabilize the lamellipodium or reinforce symmetry breaking within an existing protrusion – disrupting persistent motion (33–35). Here, we again find an effect of cell shape on protrusion stability and persistence, in which existing protrusions may either become unstable (“splitting”) or break symmetry (“angular diffusion”) at certain parameter values (Figure 4).

Given this interplay between cell shape, speed, and turning behavior, our model demonstrates how cell shape puts an upper bound on both migratory speed and persistence: both saturated at high *λ*_act_ levels where the cell had attained a broader, more keratocyte-like shape (Figure 4). A similar saturation of the shape-speed correlation at broad cell shapes was found in fish keratocytes (9). These observations clearly show that not only persistence, but also cell speed has a natural upper bound determined at least partially by cell shape dynamics.

Our results seem to suggest a role for dimensionality in this process, as the saturation of speed and persistence behaved differently in 2D versus 3D. However, we note that our cells behaved slightly differently in 3D; they had a harder time forming stable protrusions, but when they did, they almost always took on a broad, keratocyte-like shape (at least temporarily). This seemingly contradicts *in vivo* movies of T cells moving in a 3D environment such as the lymph node (16, 36), where cells *do* seem to be able to move in an elongated shape for prolonged periods of time. On the other hand, it should be noted that while cells may move freely on the empty surface of a 2D petri dish, there is no such thing as “free” migration in 3D; in reality, cells migrating in 3D always encounter barriers from their environment (be it the fibers of an extracellular matrix or surrounding cells). It is unclear whether the slightly different behavior in 3D is an artefact of the model, of the free environment, or both; comparing speed and persistence saturation in different 3D models (37, 38) may clarify this issue in the future. Nevertheless, these effects of dimensionality and environment do further stress the intricate link between changes in cell shape and the resulting motility patterns.

Together, these results shed new light on the saturation of persistence observed in the experimental speed-persistence curves: whereas persistence saturated before speed in all experimental settings considered by Maiuri et al (7), our model suggests that – depending on the cell’s shape – scenarios in which speed saturates earlier could likewise exist.

The link between cell shape and intracellular dynamics is further influenced in the CPM by the environment, which strongly constrains the shape and direction of protrusions formed. For example, in our *in silico* model of T cell migration in the epidermis, environmental constraints posed by the dense keratinocyte layer restricted persistent movement and obscured the UCSP (Figure 5), showing that environmental constraints can overrule the UCSP in at least some of the environments cells face *in vivo*. We therefore predict that speed-persistence coupling may not be visible in *in vivo* imaging data of T cells patrolling the epidermis: in such an environment, both speed and persistence likely reflect the maximum of what is feasible given the environmental constraints rather than being the result of an intrinsic coupling. In complex, highly restrictive environments, cells may choose the path of least resistance (39), with a lesser role for their intrinsic polarity mechanism in this context.

But compared to most other tissues, the epidermis is an extreme example of a confining environment. When the constraints on cell movement are less stringent, the UCSP could have a larger role in determining persistence. For example, Read et al. (14) described a correlation between cell speed and turning behavior among T cells migrating in an inflamed lymph node. Although they did not explicitly link this to the UCSP, they found that random walk models incorporating this correlation captured the data better than those that did not – suggesting the UCSP poses an essential constraint on *in vivo* motility in at least some settings.

The universality of the UCSP has important implications for computational models of cell migration in general. For example, in the last decade, several studies have investigated the functional consequences of T cell motility patterns: how should T cells move to find their targets most efficiently? Using mathematical or agent-based variations of random walk models, these studies compare different migratory strategies in terms of “search efficiency” (11, 40–43). However, selecting and fitting these models is difficult; multiple models can often fit the same experimental data depending on the metrics used to quantify migration (14, 43, 44), and even slight differences in the model used can have large consequences for the area exploration predicted on time scales beyond that of the experiment (45). Moreover, these models treat speed and persistence as input parameters that can be independently tuned, when they are in fact linked through the UCSP. This may yield models that seemingly fit the data but in truth reflect motility patterns impossible for a real cell to adopt. Indeed, two recent studies showed how the UCSP can alter cell motion patterns and space exploration (46, 47). Incorporating the UCSP into our models, or using models like the CPM in which it arises naturally, may be crucial to focus our research on those migration patterns that are actually attainable by real cells.

## CONCLUSION

An *existing* model like the Act-CPM can spontaneously capture the UCSP, even if it was not designed explicitly to do so. In general, we stress that in-depth analyses of published models, rather than only novel ones, can greatly advance our understanding of phenomena like the UCSP. This holds, for example, for other existing models in which cell shape, speed, and persistence emerge from a protrusion mechanism combined with membrane tension (48–50), and for the several variations of the original UCSP model that have now been developed (31, 51, 52). Studying the similarities and differences between these models may further clarify how cell shape and motility interactions can alter the shape of the speed-persistence curve.

## AUTHOR CONTRIBUTIONS

IW, IN, NG, RdB, and JT designed the research. IW, IN, and MK performed simulations and analyzed the data. IW and JT wrote the paper; MK, NG, and RdB critically revised the manuscript.

## ACKNOWLEDGMENTS

JT was supported by the Dutch Cancer Society - Alpe d’HuZes foundation (project 10620) and NWO (grant VI.Vidi.192.084). IW was supported by a PhD grant of the Radboudumc.

## SUPPLEMENTARY MATERIAL

Supplementary materials to this article are attached below.

## APPENDIX A: A COMPARISON WITH THE ORGINAL UCSP MODEL

We here compare (qualitatively) the parameters and behaviour of the Act-CPM to those of the original UCSP model as described by Maiuri et al. (7).

First, we note some interesting parallels between the protrusion parameters max_act_ and *λ*_act_ in our model and the parameters *C*_*s*_ and β of the Maiuri model. In general, the parameter *C*_*s*_ L/*c*_tot_ in the Maiuri model controls the front-back gradient of the polarity molecule and thus the strength of cell polarization. *C*_*s*_ describes the maximal concentration of active polarity molecules; above *C*_*s*_, increases in concentration offer no further stabilization of cell polarity. At low *C*_*s*_, this critical concentration is reached close to the cell border, so the polarization only spans a small region at the cell’s leading edge where *C* < *C*_*s*_. By contrast, a higher *C*_*s*_ allows this polarization to extend further into the cell – akin to how higher max_act_ values allow activity gradients to extend further into the Act cell. In the Maiuri model, the internal actin flow (coupled to speed) then maintains the front-back gradient by transporting the polarity cue. Something similar happens in the Act-CPM: the activity gradient causes the cell to extend at the front, while the volume and perimeter tension prevent similar protrusions at the back. This not only ensures that the cell protrudes at the front and retracts at the rear (translation, speed), but also ensures that the front keeps gaining active pixels while the rear rarely does (polarity, persistence).

A similar link exists between the *β* parameter in the Maiuri model and *λ*_act_: *β* defines the coupling strength between cell polarization and actin retrograde flow, while *λ*_act_ tunes the strength of positive feedback from the activity gradient on cell movement. Thus, where both *C*_*s*_ and max_act_ put an upper bound on the *size* polarization gradient in the cell, *β* and *λ*_act_ regulate how *strongly* this gradient affects motion.

In line with this idea, the effect of *λ*_act_ and max_act_ on cell migration patterns is qualitatively similar to that of *β* and *C*_*s*_, and the phase diagrams strongly resemble each other (compare Figure 2D to the phase diagram in (7)). At very low *β*/*λ*_act_, motion is Brownian because there is no mechanism to strengthen cell polarity once an active protrusion forms. Higher *β*/*λ*_act_ values do allow active migration in a *C*_*s*_/max_act_-dependent manner. Low *C*_*s*_/max_act_ values favor amoeboid “stop-and-go” motion, because the polarized region of the cell is very small and therefore easily destabilized by stochastic changes in cell motion. An increase of *λ*_act_ in these cells decreases the amount of time spent in stops (Appendix B) – which Maiuri et al also observed for an increase in *β* at low *C*_*s*_ (7). Increasing *λ*_act_ values also yield both a broader cell shape and higher speeds, consistent with the results Lavi et al. (31), who recently published a 2D adaptation of the original UCSP model. By contrast, high *C*_*s*_/max_act_ values allow a more stable cell polarization that extends further into the cell and does not decay, leading to persistent motility.

Although these observations provide a qualitative link between the parameters in the Maiuri and Act-CPM models, it is important to realise that the *C*_*s*_/max_act_ and *β*/*λ*_act_ parameters are not completely equivalent. For example, max_act_ and *λ*_act_ are more strongly linked to each other than *C*_*s*_ and *β*, in that cells with a high max_act_ require lower *λ*_act_ to start moving. Another important difference is the effect of *λ*_act_, which not only strengthens the activity feedback, but also weakens the effect of other constraints in the Act-CPM: when *λ*_act_ increases the weight of the activity feedback in the Hamiltonian, it automatically decreases the weight of the other terms. Very high *λ*_act_ values therefore lead to artefacts such as cell breaking, because the Hamiltonian no longer reflects the other important physical constraints on the cell. Thus, although parallels between the Act-CPM and Maiuri models do exist, the two are not equivalent.

## APPENDIX B: THE STOP-GO ENERGY BARRIER LINKS *λ*_ACT_ TO THE PERSISTENCE TIME

We here examine the phase diagram in more detail and show why persistence time in the Act-CPM depends “exponentially” on the *λ*_act_ parameter. Throughout, we will use a strongly simplified model for reasons of clarity.

## Simplified model: the one-pixel microchannel

Consider the following model of a cell in a microchannel that is one pixel high:

**Figure A1:**
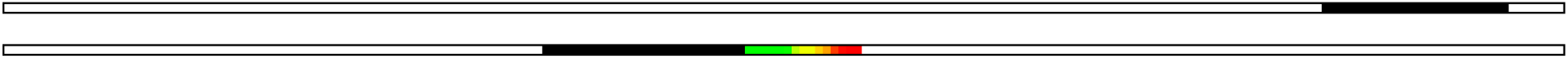
A simple model: an Act cell in a one-pixel microchannel.

The microchannel already constrains the cell shape to such an extent that we don’t need to use a perimeter constraint, so Δ*H* depends solely on adhesion, volume, and activity:

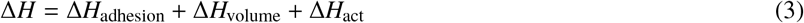

Further note that *H*_adhesion_ consists of a (fixed) contact energy *J* for each pair of neighboring pixels *(i*, *j) belonging to different cells*. A single cell only has two such pairs (*i*, *j*): at its front and at its rear. Thus, *H*_adhesion_ = 2*J* as long as there is a cell, and Δ*H*_adhesion_ = 0 for any copy attempt between two such configurations. Since the volume constraint prevents the cell from disappearing, this always applies and we can safely neglect Δ*H*_adhesion_, leaving only the latter two terms.

We now examine how these competing energy terms determine motility.

## States, phases, and the energy barrier

An Act cell in a 1D microchannel can attain two states: a non-motile “stop” and a motile “go” state. These relate to – but are different from – the “phases” in the phase diagram: an NM-cell spends all its time in *stop*, a P-RW cell spends all its time in *go*, and an I-RW cell alternates between the two (Figure A2A).

**Figure A2:**
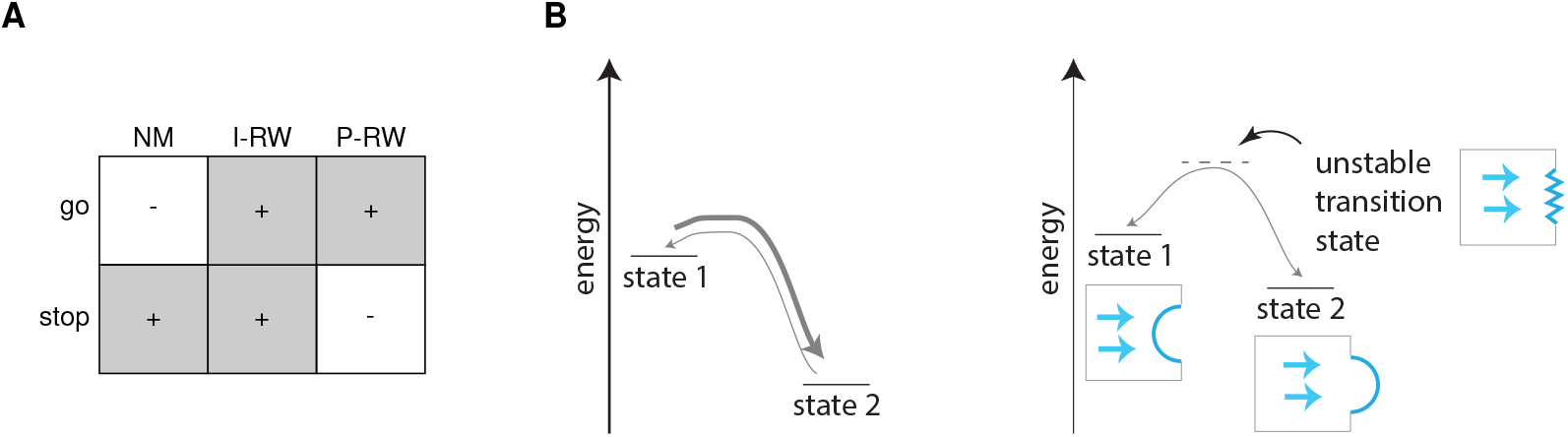
State transitions. (A) Relationship between “states” (stop, go) and “phases” (NM, I-RW, P-RW). (B) Energy diagram of a state transition. In principle, when a system can switch between two states, it will switch towards the state with lower energy at a higher rate (left). However, the switching rate also depends on the *energy barrier* that must be crossed to switch to the other state: a large barrier can prevent switching, even towards lower energy (right). The system can still switch between the two states, but does so only rarely – maintaining each state for some time before managing to cross the barrier.

But even though the I-RW cell can be in both states, it does not switch between the two continuously; it maintains each state for some time. The “stop-and-go” movement of an intermittent random walk (I-RW) is well-described as a system with two states – “stop” and “go” – for which the difference in *energy* controls transitions between them (Figure A2B).

Such state transitions are well-known from systems such as chemical equilibria, where a molecule can exist in one of two relatively stable forms and transitions between them at some rate. Each form has an energy, and additionally, the transition may require the molecule to first take on a third, “transition state” that is unstable (has a higher energy). This results in an “energy barrier” between the two states that can be crossed every once in a while, but still keeps the molecule stable in one of the two states for at least some period of time (Figure A2B).

The “stop” ↔ “go” transitions in the CPM determine motion patterns – but to understand their exact effect, we will need a more formal description of what these “states” are in the CPM, and how we can measure their energies.

## The empirical energy diagram

Of all possible grid configurations in our CPM, most fall into two basic categories: the cell is either (1) moving (i.e., it has a stable protrusion), or (2) arrested (no stable protrusion). We can define these as the “go” and “stop” states of the cell (Figure A1), which have distinct energies (explained below). The question is then: how do we measure the energy of a grid configuration, and how do configuration energies differ between the two states?

Because the CPM revolves around the global energy *H*, defining the energy of a grid configuration should be easy in principle. For example, the volume energy in our single cell simulation equals:

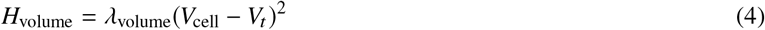

with *V*_*t*_ the target volume the cell tries to maintain. This energy depends *only* on the current configuration of the grid, which determines *V*_cell_ (the current number of pixels of our cell). The model parameters (*λ*_volume_, *V*_*t*_) remain fixed during the simulation. Δ*H*_volume_ is then simply the difference between the energies of two configurations. ^1^

Unfortunately, it is not that simple for the Act extension. Instead of specifying a *H*_act_ for each grid configuration, we now directly assign an energy *difference* Δ*H*_act_ to a specific copy attempt. This is because Δ*H*_act_ depends on *which* source pixel is trying to copy into *which* target pixel – not just on the grid configuration. To nevertheless define the energy of a cell in a particular configuration, we therefore consider the *potential energy* from having an active protrusion:

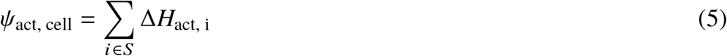

with *S* the set of all possible “protruding” copy attempts for that cell (here, that set contains just two potential copy attempts: the cell can extend either its front or its rear). The energy *H*_cell_ of a configuration is then the sum of *H* _volume,cell_ and *Ψ*_act, cell_. Tracking these energies during a simulation (Figure A3A,B), we obtain an empirical energy diagram (Figure A3C).

**Figure A3:**
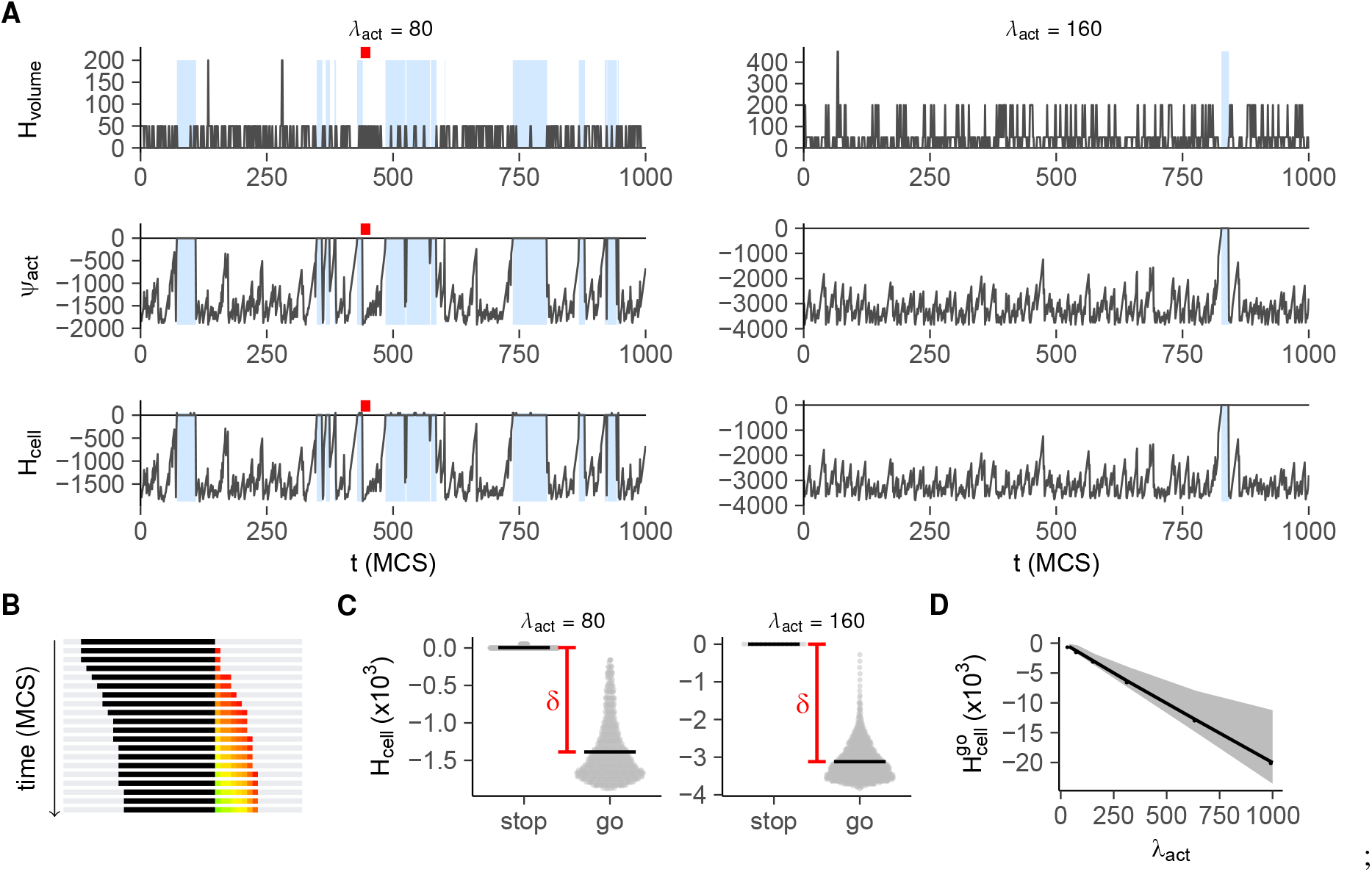
Energy diagram of the stop go state transition (*V* = 25, *λ*_*V*_ = 50,max_act_ = 25, *T* = 20). (A,B) Energies *H*_volume_, *Ψ*_act_, and *H*_cell_ during a simulation at two values of *λ*_act_. Blue shaded areas mark the “stop” state (where Δ*H*_act_ = 0), red marks highlight an interval for which the cell is shown in (B). (C) Mean and distribution of *H*_cell_ per state; each dot is one measurement (every MCS, first 1,000 MCS). (D) *H*_cell_ of the go-state at increasing *λ*_act_ values (mean±95% CI; 10,000 MCS simulation).

Here, we have defined grid configurations with *Ψ*_act,cell_ = 0 as “stop”, and all others as “go” (Figure A3A). Thus, the two states differ qualitatively: in the “stop” state, *Ψ*_act,cell_ = 0 by definition and the system is ruled solely by the volume energy – yielding a stationary cell with *H*_cell_ ≥ 0 that roughly maintains its target volume. By contrast, the large negative *Ψ*_act,cell_ of the “go” state keeps *H*_cell_ < 0, and allows the cell to extend beyond its target volume more frequently (see Figure A1 and A3A, top).

Even though 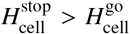, a cell in a stop does not always immediately revert to the go state because this *transition* is itself slightly unfavourable. Consider the example stop→go transition shown in Figure A3B. The cell starts in a stop (no active pixels), and then extends one pixel beyond its target volume (which is energetically unfavourable). It now has an active pixel, but the resulting grid configuration is still technically a stop: since Δ*H*_act_ of a copy attempt depends on the *geometric mean* of activities in the neighborhood of the protruding pixel, a single active pixel still yields Δ*H*_act_ = 0 and thus *Ψ*_act_ = 0. This configuration is unstable because the cell could easily lose its active pixel again to regain its target volume. The protrusion only becomes stable after a second successful copy attempt in the same direction, when *Ψ*_act_ drops below zero and enters the go-state (see also the sudden drop in energy in Figure A3A). Thus, to enter the energetically more favourable go-state, the cell must first succeed at *unfavourable* “transition” copy attempts – it must cross an energy barrier. Note, though, that this energy barrier is quite small: *H*_volume_ never becomes very high, nor does *H*_cell_.

The energy barrier is much larger for a go→stop transition, since the energy difference 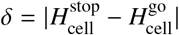 now becomes a barrier to cross. To visualize this difference, we grouped the energies from Figure A3A by state (Figure A3C). While 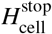 was either 0 or slightly positive, 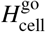 took on a range of negative values. As shown by Figure A3A, energies tended to “drift” upwards *within* the distribution of possible energies 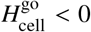 (Figure A3C) before a transition to the stop-state; effectively reducing the energy barrier for the go→stop transition. 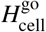 had a similarly broad distribution for different values of *λ*_act_, its mean decreasing linearly with *λ*_act_ (Figure A3D).

What does this energy diagram mean for motion patterns in our CPM? For that, we must consider how the energy difference δ between states (Figure A3C) affects the cell’s *persistence time*.

## Persistence time and the energy barrier

Formally, the *persistence time* is the time *dτ* after which a cell’s direction is no longer correlated with its previous direction. In our simplified 1D setup, this time is well-defined: it is the time needed to lose its stable protrusion on the front ^2^. With this in mind, we can view the persistence time as the time it takes the cell to cross the energy barrier from the “go” into the “stop” state (from where it can form a new protrusion in another direction). Obviously, this time will be longer if the energy barrier is steeper. From the diagram in Figure A3C, we see that the energy barrier to overcome is *at least δ* = *H*_stop_ − *H*_go_, *δ* > 0, where *H*_stop_ and *H*_go_ are the distinct energies of the stop and go states, respectively.

In analogy to the Arrhenius equation and transition state theory (53), we can view the copy success chance *P*_copy_ as a rate that decays exponentially with the energy difference Δ*H* (remember that *P*_copy_ = *e*^−Δ*H*/*T*^ for Δ*H* > 0). Now assume our state transition requires a series of copy attempts with Δ*H* > 0 adding up to *δ*:

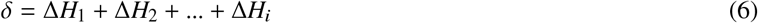

The rate with which this state transition occurs will then be:

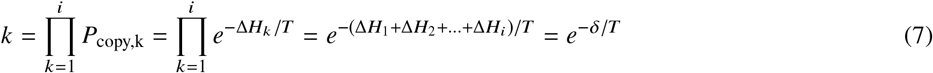

The average *waiting time* for this transition becomes:

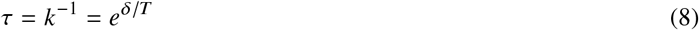

This means that the persistence time (i.e. the waiting time for a stop→go state transition) depends exponentially on the energy barrier δ. Since this energy difference depends linearly on *λ*_act_ (Figure A3C,D), we get an exponential correlation between *λ*_act_ and persistence time.

## SUPPLEMENTARY METHODS

### Simulations

Before the start of each simulation, cells were seeded in the middle of the grid and allowed a burnin time of 500 MCS to gain their optimal volume and shape. Every cell centroid was then tracked for a period of 50,000 MCS. To maximize measurement resolution while still allowing the cells to displace enough for an accurate determination of movement direction, cell centroids were recorded every 5 MCS.

### Parameters

For each experiment, we selected parameters combinations where cells had realistic shapes and migration behavior (see Table S1). Temperature, volume, perimeter, adhesion, and *λ*_V_/*λ*_P_ were chosen such that cells stayed connected even at high max_act_ and *λ*_act_ values tested (“connectedness” at least 95% for at least 95% of the time. See below). Except for max_act_ and *λ*_act_, parameters were held constant within each experiment. Parameters for 1D and 2D simulations were mostly equal – except for the larger perimeter in 1D to account for the elongated shape of cells in microchannels. For 3D simulations, we had to select other parameters to account for changes in surface to volume ratio and the thus altered relative contributions of the different terms to the total Δ*H*.

To investigate the link between speed and persistence, we analyzed cell tracks with increasing *λ*_act_ while keeping max_act_ fixed (Table S2). Max_act_ “actin lifetime” values were chosen to obtain a range of protrusion sizes. Although max_act_ can in principle take on infinitely large values, this would result in a cell where each pixel at the cell border has the same (infinitely large) activity value. In this scenario, no protrusion can form because there is no symmetry breaking and thus no cell polarization. As such behavior does not occur in real cells – that never have exactly equal actin polymerization around the entire cell rim – we did not include such large max_act_ values in our simulations. Instead, we focused on max_act_ values such that the cell went from forming small protrusions to large protrusions occupying a substantial fraction – but not all – of the cell volume (see *Analysis* of the *Methods* in the main text). For each max_act_, a range of *λ*_act_ “forces” was then chosen such that cells went from completely Brownian motion (persistence time ~ 5 MCS, the time between subsequent measurements of cell location) to maximally persistent motion (persistence time ~ 10,000 MCS). Persistence times higher than 10,000 MCS were not considered, as such high persistences will likely be underestimated due to the finite total simulation time (50,000 MCS). For skin simulations, T cell were modelled with max_act_ = 30 or 100 and variable *λ*_act_ in two different types of tissues (Table S3).

### Grid initialization

#### Microchannel simulations

To simulate migration of cells confined in a 1D microchannel, we created a 2D grid with an effective height of 10 pixels and a width of 150 pixels (but since grid borders were linked in the *x*-dimension, the effective length of the microchannel was infinite rather than only 150 pixels). Cells were confined by a layer of “barrier” pixels on the top and bottom of the grid, into and from which copy attempts were forbidden (yielding a total grid height of 12 pixels). A single cell was seeded in the middle of the channel for each simulation.

#### 2D and 3D simulations

To simulate migration in 2D and 3D, we seeded single cells in the middle of an empty 2D or 3D “infinite” grid, respectively (infinite grids were implemented as 150 × 150 or 150 × 150 × 150 grids with linked borders).

#### Skin simulations

For simulations of T cells migrating in the epidermis, 30 keratinocytes were seeded randomly on a 150 × 150 pixel “infinite” grid (with linked borders). To obtain a tissue tightly packed with intact cells, keratinocytes were initially seeded with a tighter perimeter P of 200, and each cell was given a burnin period of 50 MCS before a new cell was seeded. After an additional burnin phase of 500 MCS to let the tissue equilibrate, keratinocytes were given back their original perimeter value (Tables S1), and the cell in the middle of the grid was changed into a T cell before starting the simulations.

### Connectedness

The connectedness *C* represents the probability that two randomly chosen pixels from the same cell are part of a single, unbroken unit. To compute *C*_*i*_ of a cell *i*, we represent the cell *i* as a graph *G*_*i*_ where every node *p* is a pixel belonging to cell *i*, and pixels are connected by an edge if they are adjacent to each other on the CPM grid (that is, if they are from the same Moore neighborhood). We then group the pixels into *n connected components* [*c*_1_, …, *c_n_*] – that is, groups of pixels where for every pair of pixels (*p*_1_ ∈ *c*, *p*_2_ ∈ *c*), it is possible to walk from *p*_1_ to *p*_2_ via the edges of graph *G*_*i*_. We then define *C*_*i*_ as:

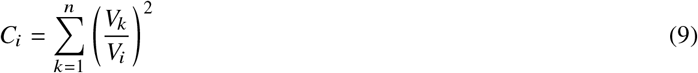

where *V*_*k*_ is the pixel volume of connected component *c*_*k*_ in *G*_*i*_, and *V*_*i*_ the total volume of cell *i*. Thus, an unbroken cell – which by definition has only one connected component – has *C*_*i*_ = 1, whereas a cell broken into many isolated pixels has *C*_*i*_ → 0.

### Phase diagram

To compute a phase diagram of migration modes, we performed microchannel simulations at max_act_ values ranging from 0 → 100 and lambda_act_ values ranging from 0 → 1500 (30 independent simulations per parameter combination). Centroids were recorded as usual and used to compute displacements Δ*x* (along the direction of the microchannel) over 10MCS.

Connectedness (see above) was tracked during the simulation; cells were considered broken if the connectedness was < 0.95 for more than 5% of the simulation. For the *N* non-broken cells, displacements were shuffled and redivided to get *N* independent displacement distributions, each consisting of displacements from different, independent cell tracks. This shuffling step is necessary for a robust estimate of the distribution of displacements a cell at that parameters may choose: a persistent cell moving to the left may only have a single peak at negative Δ*x*, yet that does not mean that another cell at the same parameters may not move to the right instead. By shuffling the displacements, we reduce the impact of interdependency of the displacements, while still retaining separate estimates to quantify uncertainty from.

The displacement distributions obtained in this way were fitted using a Gaussian mixture model to determine the number of peaks (using the R package mclust, v5.4.6 (54); using the function *Mclust* with 1-3 clusters and “modelNames” = “V”). Each distribution was then classified as follows:

- Non-migratory (NM) if the fitted mixture model contained a single peak, or if it contained two peaks but these were not clearly separated (< *σ* apart, where *σ* is the maximum of the two SDs of the mixture model). This extra filtering step was added to prevent non-motile cells with a single peak slightly deviating from a normal distribution from being classified as P-RW (which they are clearly not).
- Persistent random walk (P-RW) if the fitted model contained two clearly separated peaks (see previous point), or if it contained three peaks but the middle peak (at 0) had a mixing proportion of less than 5% (this extra filter prevents cells from being incorrectly classified as I-RW when the algorithm overfits noise with an extra peak that is not really there).
- Intermittent random walk (I-RW) if the fitted model contained three peaks, and the mixing proportion of the middle peak at zero (representing the “stops”) was <5%.
- Broken if the connectedness of the cell was below 0.95 for at least 5% of the time; see above.

This classification procedure yielded 30 independent classifications per parameter combination. For the phase diagram, we used the most frequent label as the final class, with a color intensity depending on how many of the independent estimates had this class.

### Speed and Persistence

To compute speed and persistence, we performed step-based analyses on groups of 5 simulated tracks at the same time, yielding 6 estimates of speed and persistence for every parameter combination (30 simulations total in 6 groups of 5 tracks). These values were then used to estimate the overall value (mean) and variation (standard deviation) of speed and persistence at that parameter combination. The choice to analyze tracks in groups of 5 ensures that step-based analysis (such as computing the autocovariance curve for persistence, see below) are more robust, since they contain steps from several independent tracks. At the same time, these groups are small enough that several estimates can be made from the total 30 simulations – allowing us to provide a measure of uncertainty as well.

#### Speed

To compute speeds for the speed-persistence plots, we first computed the mean step-based instantaneous speed in every track analysis group of 5 tracks (see above) using the “speed” function of celltrackR.

For the speed distribution curves, instantaneous speeds were computed over time intervals of 10 MCS and then pooled from all 30 simulations at that parameter combination. To ensure that nothing in these curves could be an artefact of cell breaking, we filtered on those 10 MCS intervals where the connectedness was exactly 1.

#### Persistence

To measure the persistence of a moving cell, consider the vectors 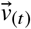 (movement direction at time *t*) and 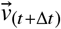 (movement direction at time *t* + Δ*t*). When the cell moves persistently, we expect that the direction of movement at *t* + Δ*t* is similar to that at *t*, even for relatively large values of Δ*t*. By contrast, for a cell undergoing random Brownian motion, the direction of 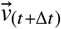 is probably unrelated to that of 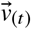.

To quantify this, consider the dot product between the movement vectors 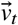 and 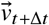:

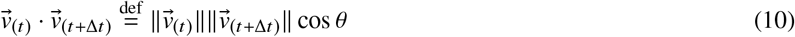

Here, cos *θ* of the angle between vectors 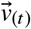 and 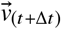 is 1 when the vectors align perfectly (*θ* = 0), −1 when they are exactly opposite (*θ* = 180), and somewhere in between for all other angles. When we take Δ*t* = 0, equation 10 simplifies to:

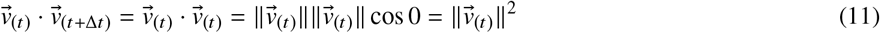

As 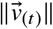 equals the instantaneous speed at time *t*, the average of this dot product for different values of *t* with Δ*t* = 0 is just the squared mean speed 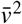.

However, when we increase Δ*t*, the vectors 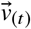 and 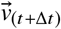 are no longer perfectly aligned, and their dot product becomes smaller. The rate at which this decay occurs depends on the motility mode of a cell: for a given Δ*t*, persistent cells will on average have a smaller θ and thus a larger dot product than cells undergoing Brownian motion. Thus, to compute persistence, we first construct the *autocovariance curve* of the average dot product 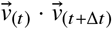 along a track as a function of Δ*t* (using the “overallDot” function of the MotilityLab package). As a measure of persistence, we then compute the half-life τ of this autocovariance curve for which:

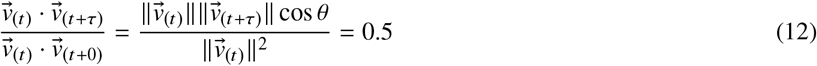

As the dot product decays more slowly for more persistent cells, high values of τ indicate persistent movement. Note that, as the 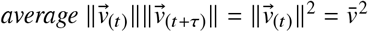, τ is independent of the mean speed 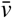, even though the dot product is not.

Technical note: for very high persistences, it can be difficult to estimate the autocovariance half-life. If the autocovariance curve fails to drop to sufficiently low values within the time scale of the simulation, it becomes impossible to reliably estimate its decay. To avoid this problem, we only compute persistences for those groups of tracks where the autocovariance curve dropped to <10% of its initial value, returning NA otherwise. The duration of the simulation (50,000 MCS) was sufficient that this never happened even at the highest persistences reported in this paper.

## SUPPLEMENTARY TABLES

**Table S1:**
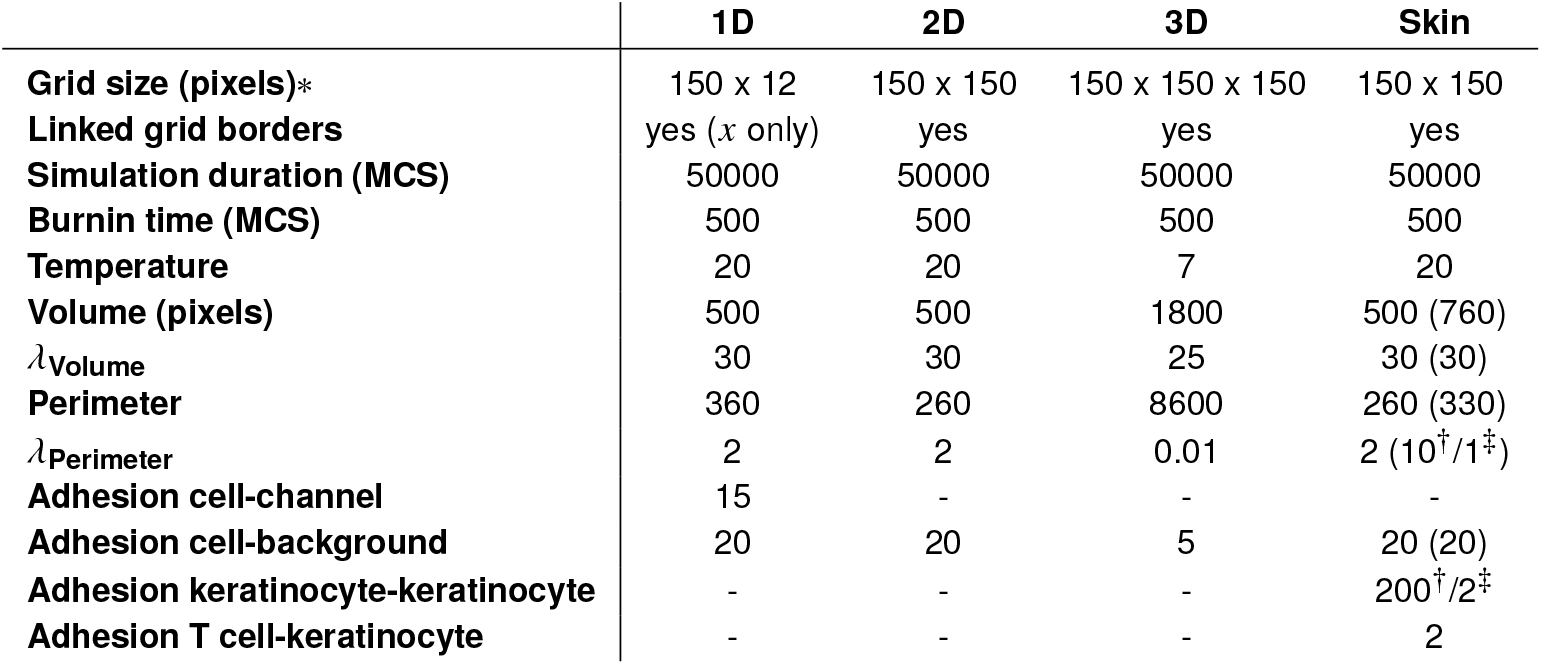
CPM parameters used in different experiments. Within each experiment, parameters were kept constant. Skin simulation parameters apply to the T cells; bracketed parameters indicate values used for the keratinocytes. While these grid dimensions are finite, the linked grid borders (Artistoo setting: `torus=TRUE`) ensure that the effective grid size is infinite. While the grid height for microchannel simulations was 12 pixels, the upper and lower rows of pixels consisted of the microchannel walls - yielding an effective height of only 10 pixels. ^†^Value used for stiff tissue, ^‡^value used for deformable tissue.

|  | 1D | 2D | 3D | Skin |
| --- | --- | --- | --- | --- |
| **Grid size (pixels)\*** | 150 x 12 | 150 x 150 | 150 x 150 x 150 | 150 x 150 |
| **Linked grid borders** | yes (x only) | yes | yes | yes |
| **Simulation duration (MCS)** | 50000 | 50000 | 50000 | 50000 |
| **Burnin time (MCS)** | 500 | 500 | 500 | 500 |
| **Temperature** | 20 | 20 | 7 | 20 |
| **Volume (pixels)** | 500 | 500 | 1800 | 500 (760) |
| $\lambda_{\text{Volume}}$ | 30 | 30 | 25 | 30 (30) |
| **Perimeter** | 360 | 260 | 8600 | 260 (330) |
| $\lambda_{\text{Perimeter}}$ | 2 | 2 | 0.01 | 2 ($10^{\dagger}/1^{\ddagger}$) |
| **Adhesion cell-channel** | 15 | - | - | - |
| **Adhesion cell-background** | 20 | 20 | 5 | 20 (20) |
| **Adhesion keratinocyte-keratinocyte** | - | - | - | 200<sup>†</sup>/2<sup>‡</sup> |
| **Adhesion T cell-keratinocyte** | - | - | - | 2 |

**Table S2:** Combinations of max_act_ and *λ*_act_ used in different experiments. For each max_act_, *λ*_act_ values were chosen such that cells went from no persistence to maximal persistence (see sections and for details on parameter selection).

| Experiment | $\max_{\text{act}}$ | $\lambda_{\text{act}}$ |
| --- | --- | --- |
| 1D | 30 | 100, 200, 600, 800, 1000, 1200, 1400 |
|  | 40 | 60, 80, 100, 200, 400, 600 |
|  | 50 | 50, 60, 70, 80, 90, 95, 100 |
|  | 60 | 40, 45, 50, 55, 60, 65, 70 |
|  | 80 | 25, 30, 35, 38, 40, 42, 45 |
|  | 100 | 30, 32, 34, 36, 38, 40 |
| 2D | 30 | 50, 100, 200, 300, 400, 500, 700, 1000 |
|  | 40 | 50, 75, 100, 150, 200, 250, 300, 500, 800, 1000 |
|  | 50 | 50, 75, 100, 125, 150, 200, 300, 500 |
|  | 60 | 40, 60, 80, 100, 120, 150, 250 |
|  | 80 | 30, 40, 50, 60, 70, 100, 150, 200 |
|  | 100 | 30, 35, 40, 45, 50, 60, 70, 100, 150, 200 |
| 3D | 30 | 40, 45, 47, 48, 49, 50, 52, 55, 60 |
|  | 40 | 30, 35, 37, 38, 39, 40, 42, 45, 50 |
|  | 50 | 25, 30, 33, 34, 35, 36, 37, 38, 40 |

**Table S3:** Combinations of max_act_ and *λ*_act_ used in the skin simulations. For all combinations, simulations were performed both in “stiff” and “deformable” tissue (see Table S1 for the corresponding keratinocyte adhesion and *λ*_Perimeter_ values).

| Experiment | $\max_{\text{act}}$ | $\lambda_{\text{act}}$ |
| --- | --- | --- |
| skin | 30 | 200, 500, 1000, 1250, 1500, 1750, 2000, 2500 |
|  | 100 | 100, 200, 300, 500, 750, 1000 |

## SUPPLEMENTARY MOVIES

Supplementary movies are available on: https://computational-immunology.org/inge/ucsp-cpm/supplements/.

Movie S1: Act cells in microchannels have variable motility depending on their protrusion size. The movie first shows an Act cell with a small protrusion – giving rise to typical “stop-and-go” motility, followed by an Act cell with a more stable protrusion – yielding more continuous movement.

Movie S2: Act cells in 2D and 3D have both different shapes and migration patterns. The movie shows examples of typical migration patterns in 2D and 3D. At low max_act_ values, small, unstable protrusions form and decay dynamically – giving rise to amoeboid “stop-and-go” migration. At higher max_act_ values, the cell becomes broader and forms a large, stable protrusion perpendicular to its direction of movement – yielding a movement pattern similar to that observed in fish keratocytes (“keratocyte-like”).

Movie S3: Different processes put an upper bound on persistence depending on the cell shape. For cells with an unstable protrusion (low max_act_), active protrusions can split into two parts whenever the cell tries to form protrusions that are too broad to maintain. This forces the cell to turn in the direction of one of the protrusion halves, limiting persistence. The movie shows examples of such “protrusion splitting” (indicated by pauses and arrows). By contrast, for cells with a stable protrusion (high max_act_), broad protrusions do not split or decay – but the most active part of the protrusion can still shift away from the protrusion center, causing the cell to turn (“angular diffusion”). The movie indicates this type of stochastic turning by showing in red the movement of the cell’s center of mass.

Movie S4: Act T cells in the epidermis move by squeezing in between keratinocytes. At low *λ*_act_ and max_act_, they follow the typical “stop-and-go” motion of an amoeboid cell. At high max_act_ and/or *λ*_act_, they keep their elongated form in between the keratinocytes, rather than taking on the broad keratocyte-like shape that free Act cells would. However, they stop less frequently than cells with low max_act_ and *λ*_act_, and actively probe their environment by letting parts of their active protrusion extend into the space between keratinocytes.

Movie S5: Tissue stiffness affects Act T cell migration mode. Cells with high max_act_ are mostly prevented from broad protrusions in a stiff tissue. Their protrusions only broaden in the space between keratinocytes before rapidly splitting in two. By contrast, the same cells can form and maintain broad protrusions in a more deformable tissue by pushing apart the cells surrounding them.

## SUPPLEMENTARY SIMULATIONS

Supplementary simulations are available on: https://computational-immunology.org/inge/ucsp-cpm/supplements/.

Interactive Simulation S1: Simulation of Act cell confined between two microchannel walls. Adjust the values of max_act_ and *λ*_act_ to see their effect on cell shape and motility.

Interactive Simulation S2: Simulation of an Act cell moving freely on a 2D surface. Adjust the values of max_act_ and *λ*_act_ to see their effect on cell shape and motility.

Interactive Simulation S3: Simulation of an Act cell moving freely in a 3D open space. Adjust the values of max_act_ and *λ*_act_ to see their effect on cell shape and motility.

Interactive Simulation S4: Simulation of an Act cell moving in a tissue of densely packed keratinocytes. Adjust the values of max_act_ and *λ*_act_ to see their effect on cell shape and motility, and compare this between the “stiff” and “deformable” tissues.

## SUPPLEMENTARY FIGURES

**Figure S1:**
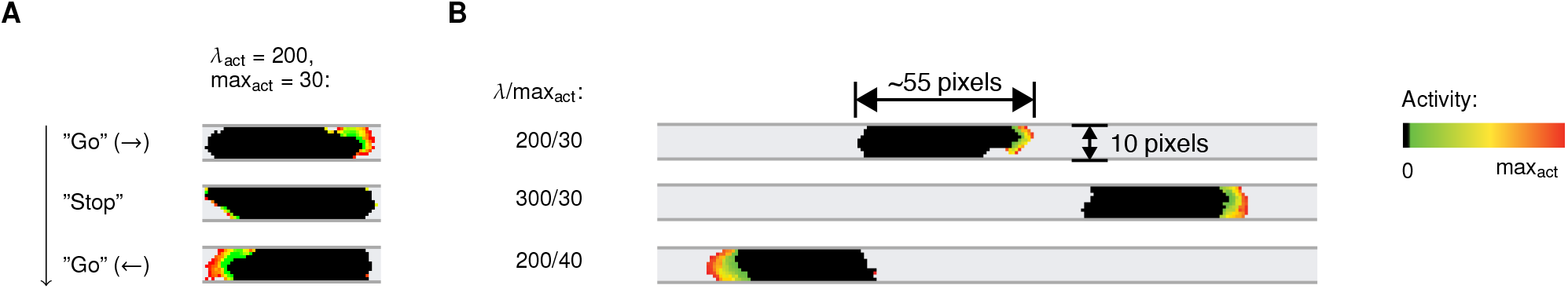
Act cells in microchannels have “stop-and-go” motility and variable protrusion sizes. **(A)** Example of “stop-and-go” motility. When a protrusion decays, the cell stops until a new protrusion forms – which may be in another direction. **(B)** Example cells for different combinations of the parameters *λ*_act_/max_act_.

**Figure S2:**
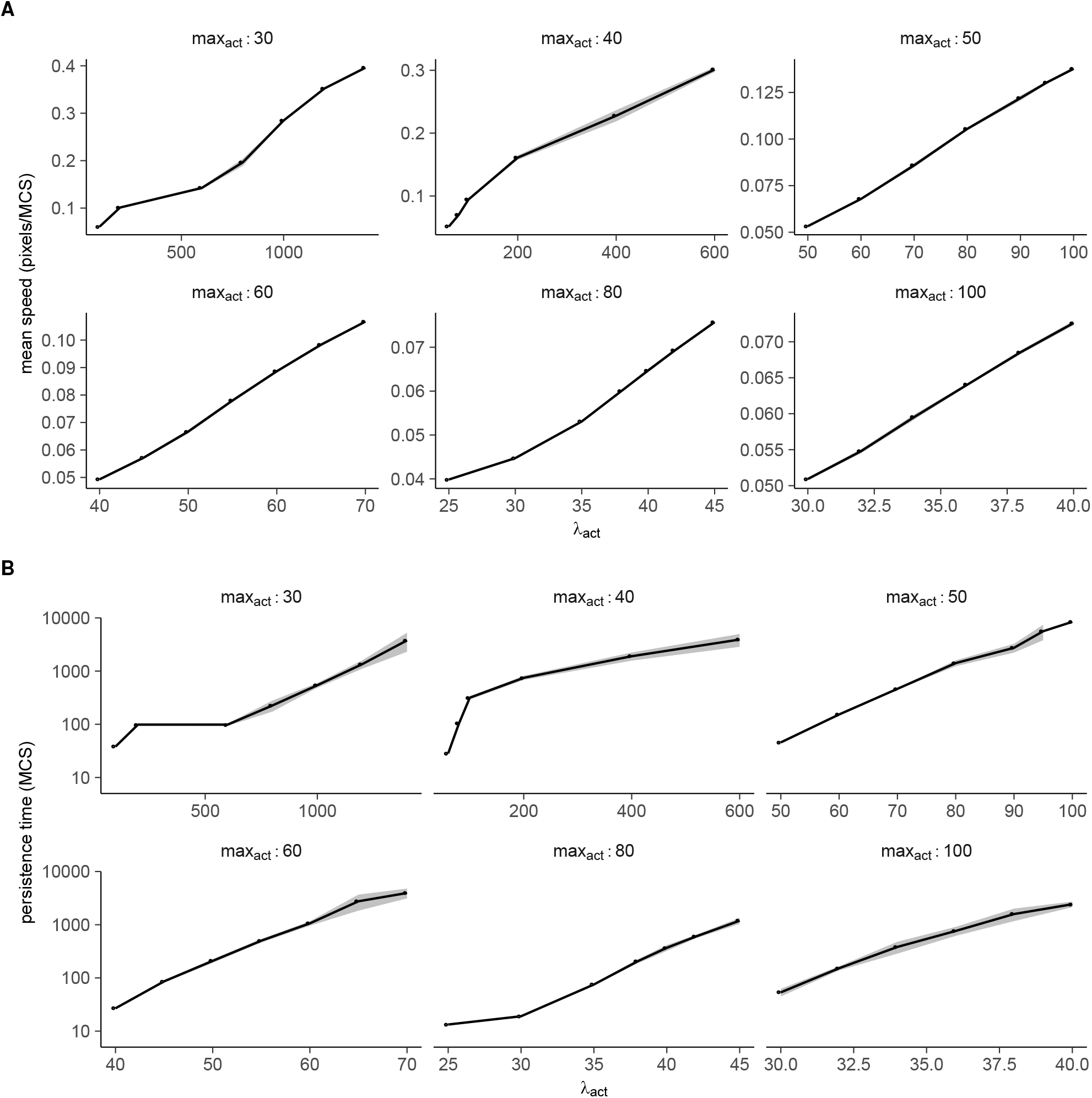
Speed and persistence depend on *λ*_act_ in a linear and exponential manner, respectively. Plots show mean ± SD of **(A)** cell speed, and **(B)** persistence as a function of *λ*_act_, for different values of max_act_. Open circles indicate points where the persistence time is lower than the time it takes the cell to move 10% of its length (corresponding to points in the gray region in Figure 2).

**Figure S3:**
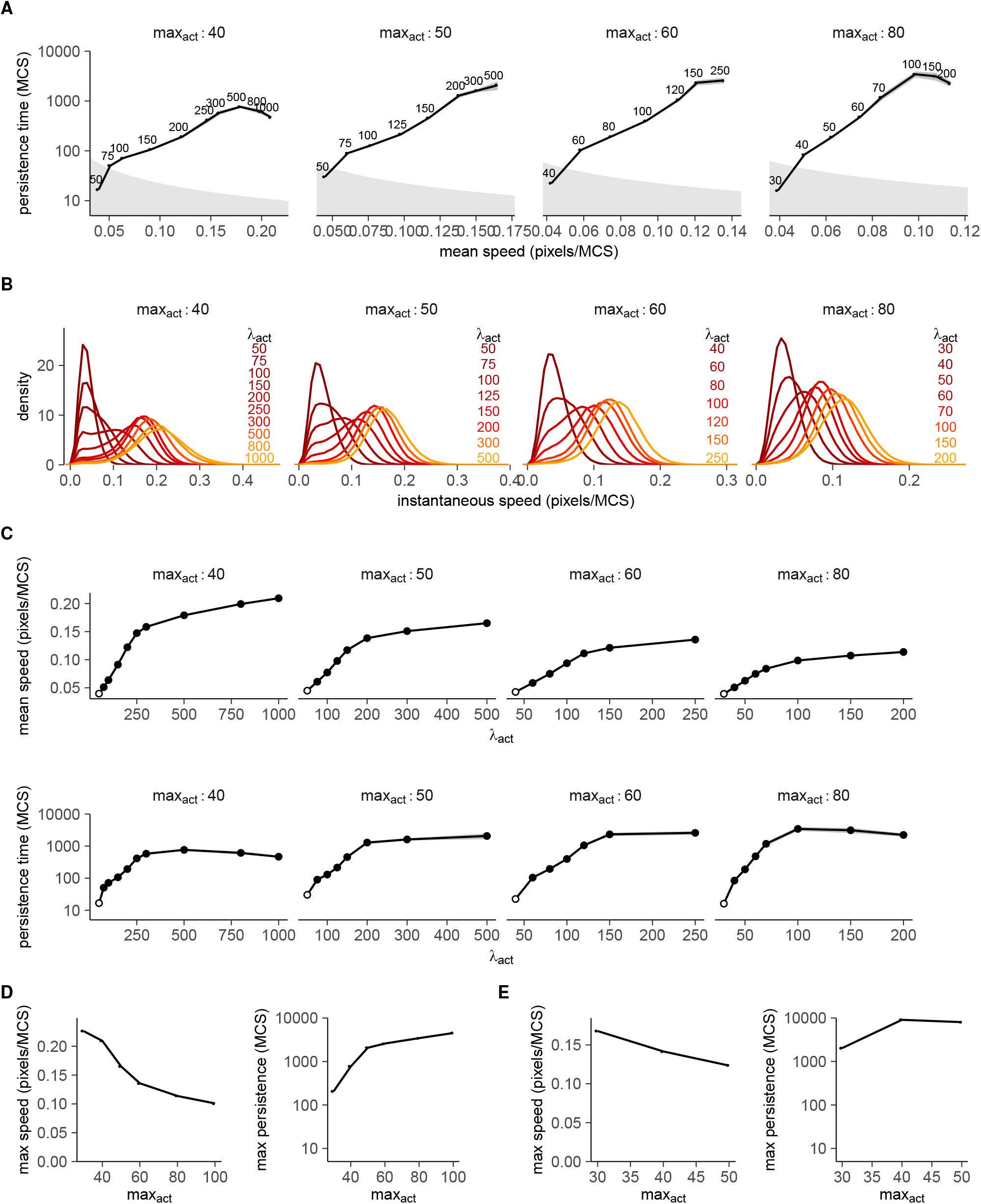
Act cells show similar behavior for different values of max_act_, see also Figure 3B,C and 4A. Plots **(A-D)** show for 2D Act cells: **(A)** Exponential speed-persistence coupling, **(B)** Distributions of instantaneous speeds, **(C)** Saturation of speed and persistence at high *λ*_act_ values, and **(D)** Maximal speed and persistence measured for all values of max_act_ in Figure 4 and panel D. **(E)** Maximal speed and persistence measured for 3D Act cells for different values of max_act_.

**Figure S4:**
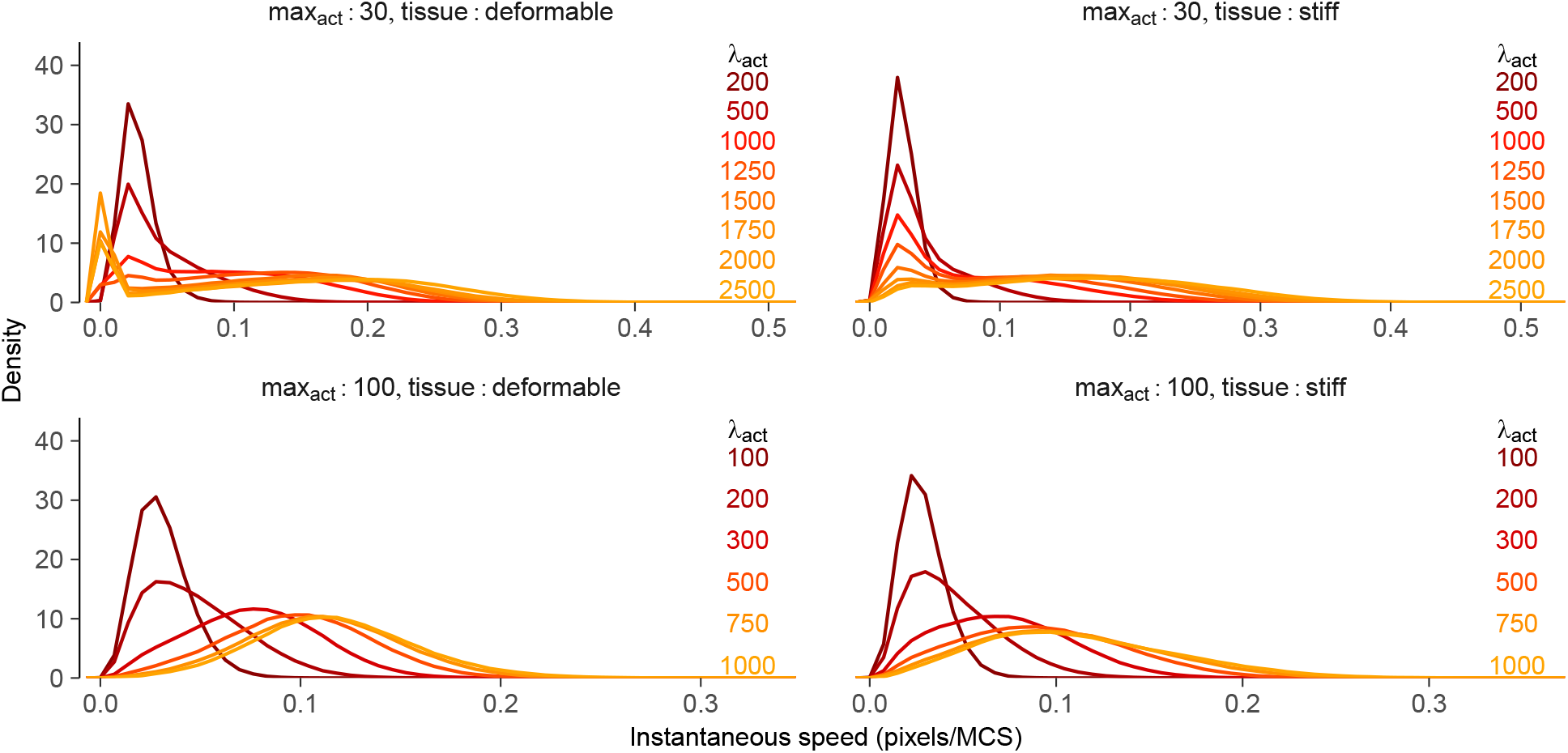
Act T cells in skin display “stop-and-go” behavior or continuous movement depending on the value of max_act_. Plots show distributions of instantaneous speed for different values of the migration parameters max_act_ and *λ*_act_, and in two tissues of different rigidity (see also Figures 3C and S3B).

**Figure S5:**
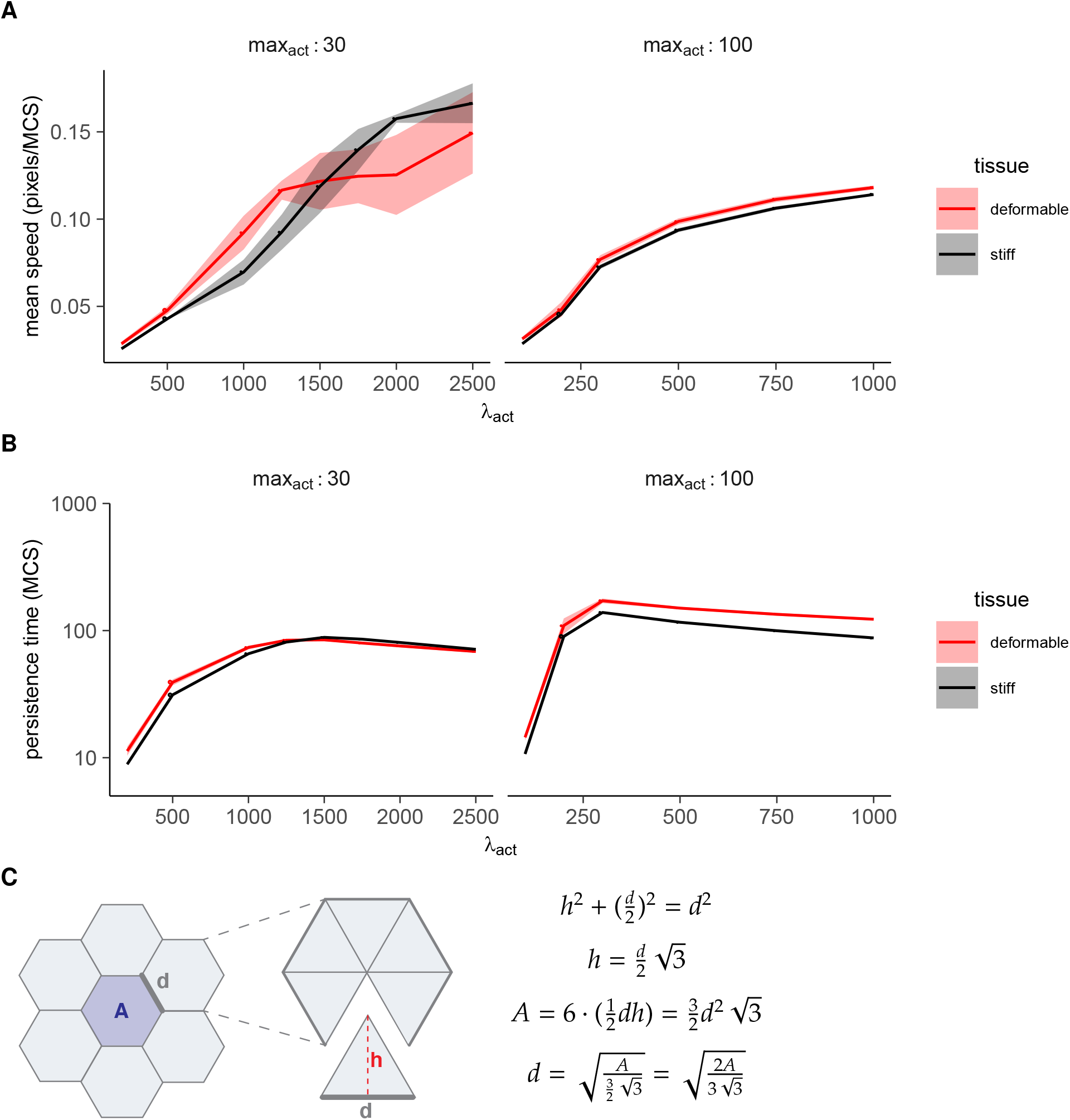
Both speed and persistence saturate for T cells moving in skin. Plots show mean ± SD of **(A)** cell speed, and **(B)** persistence as a function of *λ*_act_, for different values of max_act_ and in two different tissues. Open circles indicate points where the persistence time is lower than the time it takes the cell to move 10% of its length (corresponding to points in the gray region in Figure 5D). **(C)** If we model the keratinocyte layer as a grid of packed hexagons, the distance *d* a cell can travel before hitting a junction equals 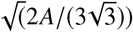 with *A* the area of the keratinocyte hexagon. Given that *A* = 760 pixels (the keratinocyte “volume” in the CPM volume constraint), we get *d* ~ 17 pixels.

## Footnotes

1 The same principle would apply to *H*_adhesion_, *H*_perimeter_ if we would use those.

2 In higher dimensions, this no longer holds because the cell can also change direction by turning, rather than losing, its protrusion. We therefore focus on the simple 1D case here.

